# A phylogenetic contribution to understanding the panzootic spread of African swine fever: from the global to the local scale

**DOI:** 10.1101/2025.08.26.672323

**Authors:** Gianluigi Rossi, E. Carol McWilliam Leitch, Jake Graham, Roberta Biccheri, Carmen Iscaro, Claudia Torresi, Samantha J. Lycett, Francesco Feliziani, Monica Giammarioli

## Abstract

African Swine Fever virus has become a primary concern for veterinarian health agencies and pig producers worldwide. The current panzootic of the virus genotype II is having a devastating impact on pig production in Africa, Europe, Asia, Oceania and Hispaniola (Caribbean). Due to its high persistence and mortality rate, disease control policies require enhanced passive surveillance activities, wild boar depopulation, containment and other costly interventions, as a safe and effective vaccine is not currently available.

Since 2007, several disease clusters have emerged far from both its original range, in South-Eastern Africa, and far from other affected suid populations. These transmissions were likely caused by anthropogenic movement of contaminated material, facilitated by the persistence of the virus in the environment and on contaminated material.

The objective of this research is to understand the spatio-temporal dynamic of the African Swine Fever virus panzootic, with a specific focus on clusters from mainland Italy. We mapped and analysed the virus spread using 228 whole-genome sequences available from online repositories and from the Italian cases/outbreaks, combined with their metadata. We inferred the pathogen phylogenies using a Bayesian phylodynamic model, with which we obtained a time-scaled and spatially explicit maximum clade credibility tree.

Our results showed a non-negligible number of long-distance transmissions across regions or continents, coupled with dense local dynamics, particularly in areas where the disease affected a naïve population. The distribution of spatial distances inferred along the trees’ branches further highlighted these trends, and revealed how previously observed survival times in pork products could allow the virus to traverse distances up to 900km. Finally, from the available sequences, we identified at least seven separate introduction events in Europe, of which at least three caused new clusters on mainland Italy.

This study provides important insights on the African Swine Fever virus introduction into many affected areas worldwide and highlights the crucial role of genomic surveillance to correctly track the pathogen spread, and to monitor the virus potential evolution.

## Introduction

African Swine Fever virus (ASFV) is one of the most serious threats to the global pig industry. The current panzootic is causing major economic losses worldwide, including in China [1], and the European Union [2], the two World’s leading pigmeat producers and exporters. In South-East and Pacific Asia, ASFV is also decimating the population of native suids, threating the food security of many rural communities [3].

The pathogen is a large (170-193 kbp) double-stranded DNA virus, first identified in Eastern Africa in 1921 [4], where it is maintained through a sylvatic cycle involving warthogs, mostly asymptomatic hosts, and soft ticks of the *Ornithodoros* genus [5]. The disease causes a haemorrhagic fever in domestic pigs and wild boars, with virulent strains that can lead to up to 100% mortality in naïve populations [5]. Currently, a safe and effective vaccine does not exist [6], thus control strategies rely on mechanically preventing transmission (e.g. fencing, biosecurity, control zones) and culling [7,8]. One concerning characteristic of ASFV is its ability to survive in the environment and in uncooked pigmeat products, such as cured meat [9]. Consequently, movement of contaminated material can lead to new clusters in areas outside its original range.

The current ASF panzootic is caused by the genotype II: it was first identified in Georgia and in other Caucasus countries in 2007, including several Russian republics [10]. From the Trans-Caucasian region, ASFV spread to Eastern Europe, reaching the European Union in the Baltic countries and Poland in 2014, then to Central Europe and to the Balkan region. It further spread as far as Belgium (2018), Italy (2022), Sweden (2023) and Albania (2024) [11]. In 2018 it emerged in China [12], and between 2018 and 2019 in other East Asian countries, including Mongolia, Vietnam, Cambodia, Hong Kong, North Korea, South Korea, Laos, the Philippines, Myanmar, Indonesia, and Timor Leste [13,14]. In 2021 it was detected on the island of Hispaniola, in the Caribbean (Haiti and Dominican Republic) [15]. In many of the affected regions ASFV successfully established in the local wild boar population, which acts as a reservoir for spillover to domestic pigs [16].

This global spread is likely caused by multiple transmission routes, which allow the virus to be spread at different spatial scales. Vector mediated and animal-to-animal transmissions, either between domestic pigs, wild boars or between the two, can spread the disease at the local spatial scale. However, anthropogenic-mediated transmission such as fomites on farm visitors’ clothing, equipment, or vehicles can reach wider areas in a short time span [17]. Finally, ASF contaminated food, animal feed or other material can spread the virus at the intercontinental scale [18].

In Italy, ASFV genotype II was detected for the first time in a wild boar carcass in early January 2022, in the North-Western province of Alessandria (Piemonte region) [19, 20]. Since then, the cluster has expanded to include another four regions (Liguria, Lombardia, Emilia Romagna and Toscana), while another three clusters have emerged in non-contiguous regions: one in Lazio, within the Rome municipality (2022), one in Calabria (2023) and finally one in Campania (2023). The Lazio cluster was resolved in January 2025, whereas the others are currently on-going. From previous genome analyses [20–22], it emerged that the strains sampled in some of these clusters belong to different genetic groups, so questions were raised about the virus introduction pathway in the country.

Here, we aim to untangle the spatio-temporal dynamics of the ASFV genotype II spread by using a phylogenetic modelling approach, which exploits the mutations in the viral genome, coupled with the isolates’ metadata, to obtain georeferenced time-scaled phylogenies. We used an alignment that included 146 whole-genome sequences (WGS) deposited in GenBank [23], and 82 sequences isolated across the four Italian mainland clusters. We first analysed the global spread of ASF to understand the outbreaks detection timeline. We then focused on the four Italian clusters and their potential links with other ASFV strains detected in Europe and Asia. Our results unveiled the global dispersal dynamic of the virus genotype II and shed light on the estimated transitions between different geographical regions. Finally, we have demonstrated that the Italian clusters are probably the result of multiple introductions of viral strains from elsewhere in Eurasia.

## Material and methods

### Alignment assembling and data preparation

A total of 228 ASF virus genotype II whole virus whole-genome sequences (WGSs) were collected for the analyses. We sourced 146 WGSs from the GenBank online repository, representing most of the affected continents with available WGSs up to January 2024 [23]. Another 82 were obtained from strains sampled in the four mainland Italian clusters: from the North-West (samples from Piedmont and Liguria, the first affected regions), Rome municipality, Campania and Calabria (see Table S1 for the full list of WGSs). The software MAFFT v7.511 [24] was used to perform a multiple sequence alignment.

To run the analyses the ASFV WGSs we used the associated metadata to the best of our knowledge, namely: sample date, host (wild boar or domestic pig) and geographical location. For the latter we used the available geographic coordinates, and a macro region assigned according to the United Nations Geoscheme (see Supplementary Material for more information). The WGSs location and region is reported in Figure 1.

**Figure 1.**
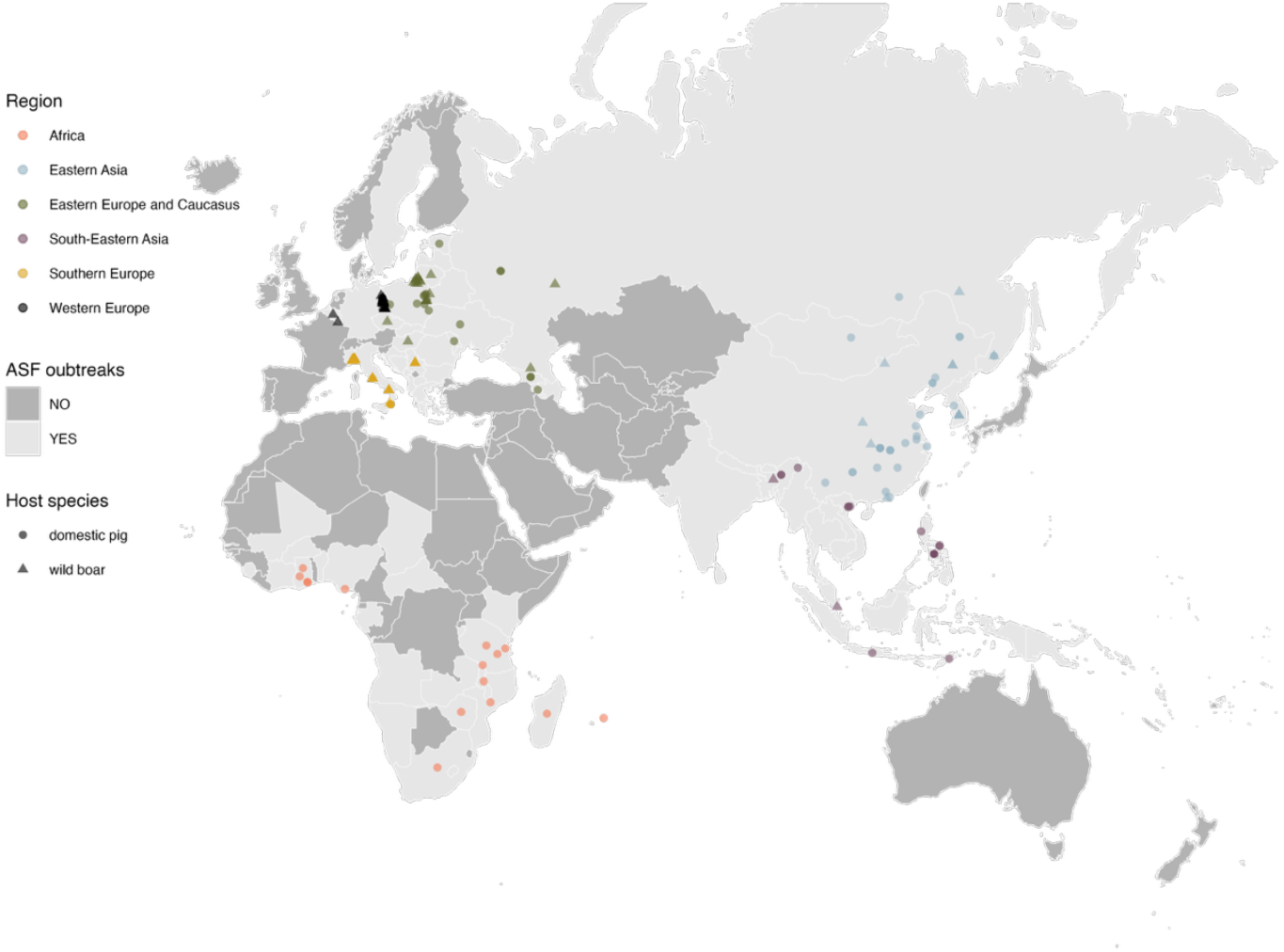
The locations of the 228 African Swine Fever virus (ASFV) whole-genome sequences used in this study. Countries coloured in light grey have recorded ASFV clusters or cases since 2007 (data from WOAH[11]). Haiti and the Dominican Republic (Caribbean) are the only countries with reported clusters of ASFV not shown on the map. The sequences’ locations are represented by dots for domestic pigs, or by triangles for wild boar (or feral pigs), and they are coloured depending on the assigned region (Africa, red; Eastern Asia, blue; South-Eastern Asia, purple; Eastern Europe and Caucasus, green; Southern Europe, yellow; Western Europe, black).

### Time-scaled trees and traits

To obtain a time scaled phylogeny and the highest clade credibility tree, we used the Bayesian Evolutionary Analysis Sampling Trees (*BEAST*) v1.10.5 [25]. The *BEAST* software uses Markov Chain Monte Carlo (MCMC) algorithms to infer values taken by multiple evolutionary parameters, such as divergence times and evolutionary rates, by repeated sampling of probability distributions. Several preliminary models in *BEAST* were run, to select the best evolutionary model (see Supplementary material). Once the best phylogenetic model was identified, the model was run again using alternative chain lengths and random number seeds to improve variability.

The obtained posterior tree samples were used as input for a further run in *BEAST* to generate a new trees distribution with trait partitions: two discrete traits, identifying the isolates region (Table 1) and host (wild or domestic), and one continuous (geographical coordinates). The chain length was 10^8^, sampled every 10^4^ to obtain a final 10,001 trees. We focused our analysis on the 9,001, after discarding 1,000 as burn-in. The maximum clade credibility (MCC) tree was extracted with *TreeAnnotator* v1.10.4, and visualised with *FigTree v1*.*4*.*4* [26], while the results were plotted in *R v4*.*4*.*1* [27].

**Table 1.** Summary of the number of ASF whole-genome sequences included in the study, divided by region and host. Armenia and Estonia had one sequence each, but they were excluded after the quality assessment.

| Region | Host | # total sequences | # selected sequences | Min year | Max year | Represented countries |
| --- | --- | --- | --- | --- | --- | --- |
| Africa | Domestic pig | 15 | 15 | 1998 | 2022 | Ghana, Madagascar, Malawi, Mauritius, Mozambique, Nigeria, South Africa, Tanzania, Zimbabwe |
| Eastern Asia | Domestic pig | 30 | 25 | 2018 | 2022 | China, Mongolia, South Korea, East Russia |
|  | Wild boar | 14 | 14 | 2018 | 2020 |  |
| South-Eastern Asia | Domestic pig | 21 | 21 | 2019 | 2023 | India, Indonesia, Philippines, Singapore, Timor-Leste, Viet Nam |
|  | Wild boar | 2 | 2 | 2020 | 2023 |  |
| Eastern Europe and Caucasus | Domestic pig | 16 | 12 | 2007 | 2019 | Czech Republic, Georgia, Hungary, Lithuania, Moldova, Poland, West Russia, Ukraine (Armenia, Estonia) |
|  | Wild boar | 18 | 18 | 2014 | 2019 |  |
| Southern Europe | Domestic pig | 12 | 11 | 2022 | 2023 | Italy, Serbia |
|  | Wild boar | 75 | 74 | 2019 | 2023 |  |
| Western Europe | Domestic pig | 2 | 2 | 2021 | 2021 | Belgium, Germany |
|  | Wild boar | 23 | 23 | 2018 | 2021 |  |

### Branches distance analysis

The model described above allows *BEAST* to estimate the coordinates of all internal nodes, and therefore we can obtain an estimate of each branch distance across the 9,001 trees. However, because estimates are done in continuous space, internal nodes locations might fall in areas with no disease observation or, in some cases, areas lacking a host species. To overcome this issue, for each of the 9,001 tree we identified which internal node was initially estimated in non-affected areas, and we rewired the branch to its parent. We continued to do so until we found a parent in an ASFV affected country, defined following the cases data published on FAO’s Empres-I [28]. Once the rewiring was done, we recalculated all the branches distance across the 9,001 trees. We also assessed the branches according to their time difference, i.e. the time between the node and the tip estimated or observed time. We divided the sample according to few different time length thresholds, chosen according to the virus persistence in different substrates: up to 15 days, approximately corresponding to the half-life of ASFV under shipping condition (14.2 days in [29]), 102 days (15 weeks), corresponding to the lifespan of the virus in refrigerated meat [7] and 137 days, corresponding to the time a number of cured meat product were observed negative [9], and finally a year.

### Single gene analysis

We run a further analysis on three single genes that we extracted from the alignment, and we assessed the similarity of those across the alignment. We retrieved the B602L (1,593 bp), I73R (219 bp), and MGF 360-10L (1,038 bp) gene sequences from Georgia 2007/1 reference sequence on GenBank (FR682468.2, [30]). All the three genes are considered among the genetic or sub-genotype markers [31,32]. B602L is located in the virus genome central variable region, and it can discriminate strains that are identical according to their p72 and p54 genotypes [33], while I73R was found to be critical for ASFV pathogenesis as it suppresses the host innate response [34]. Finally, MGF360-10L promotes the survival in host cells and might be responsible for immune response inhibition [35,36]. We used the BLAST algorithm [37] to extract the correspondent genome sections from the alignment, and we built a phylogenetic tree for each gene using *IQ-TREE2[38]*.

## Results

### Time scaled trees and global spread

After a preliminary sequence selection phase (Supplementary material), the final alignment we used for the analyses included 217 WGSs. The substitution model that best performed in our selection process was the Hasegawa-Kishino-Yano (HKY) with Gamma-distributed heterogeneity model, coupled with exponentially distributed relaxed clock rate and the Bayesian SkyGrid[39] tree prior. The average clock rate estimate was 1.07×10^-5^ substitutions/site/year [95^th^ High-posterior density, HPD, 0.82×10^-5^-1.36×10^-5^], which correspond to 1.97 substitutions/genome/year [HPD 1.51-2.51]. We did not observe any statistically significant variation in the clock rate estimates across region or host (Figure S2). The estimated effective population size is reported in Figure S3.

In Figure 2 we reported the MCC tree with the addition of traits, and branches coloured by region (see Figure S4 and S5 panel A for the branches coloured according to, respectively, the branches posterior probability estimate and host). According to the model estimates, the most recent common ancestor (MRCA) was in October 1993 [HPD February 1984 – November 1998]. The MRCA of the Eurasian epidemic median estimate was November 2005 [HPD September 2003 – May 2007], while the MRCA for the East Asian clade was December 2016 [HPD November 2015 – September 2017].

**Figure 2.**
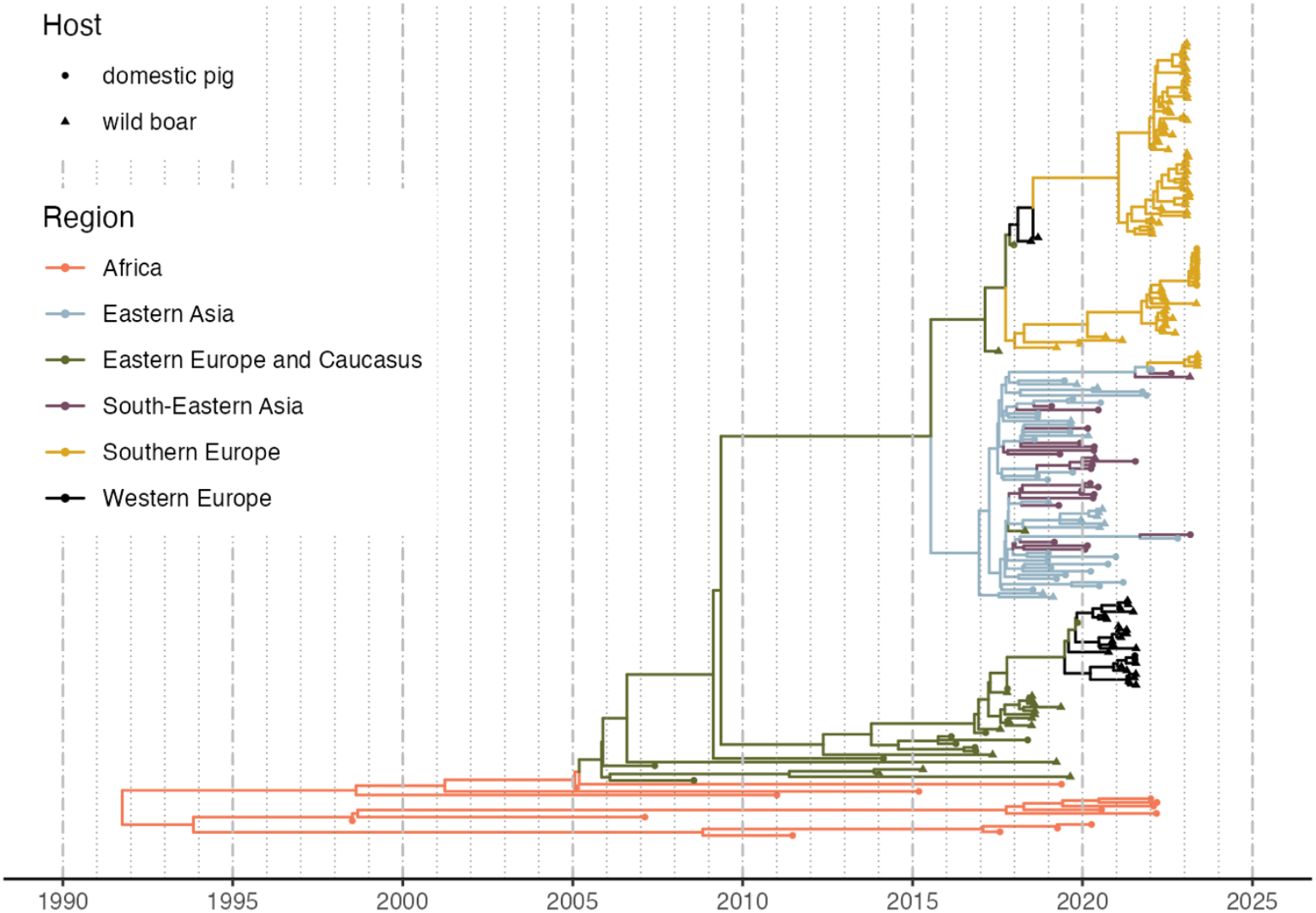
The Maximum Clade Credibility (MCC) tree of the 217 African Swine fever virus whole-genome sequences used in the study. Branches are coloured according to the most likely region, while internal nodes and tips shape are determined by the observed (tips) or inferred (internal nodes) host (dots for domestic pigs, or by triangles for wild boar or feral pigs).

The transmission matrix (Figure 3) shows the number of estimated ASFV transitions within and between regions. The estimated number of within-region transitions is highly correlated to the number of sequences from each region (Spearman ρ = 0.94). As expected, the model estimated one transition from Africa to Eastern Europe and Caucasus. Most of the other between-region transitions consist of single events (estimated transitions value: ∼ 1), with two exceptions: first, from Eastern to Western Europe (2.6). This may be a result of the introductions into Easter Germany from the contiguous areas of Western Poland, which can be considered one cross-border cluster, with the strains also showing a characteristic tandem repeat in the O174L gene [40]. The highest number of between-region transitions was observed from Eastern to South-Eastern Asia (12.2), which is of a similar magnitude to the South-Eastern Asia within-region transitions (18.2): this might have been facilitated by the trading links within countries of the area. Figure S5 reports the transition matrix between hosts (panel B): while between-host transitions are not negligible, our results show a dominant circulation within each host. The insight we can gather from this might be limited, however, because the internal nodes assignment of this trait could be easily biased by the available observations. By adding the coordinates as a continuous trait, BEAST estimated the global spatio-temporal spread of ASFV, which is reported in Figure 4 (and a dynamic movie in Movie S1).

**Figure 3.**
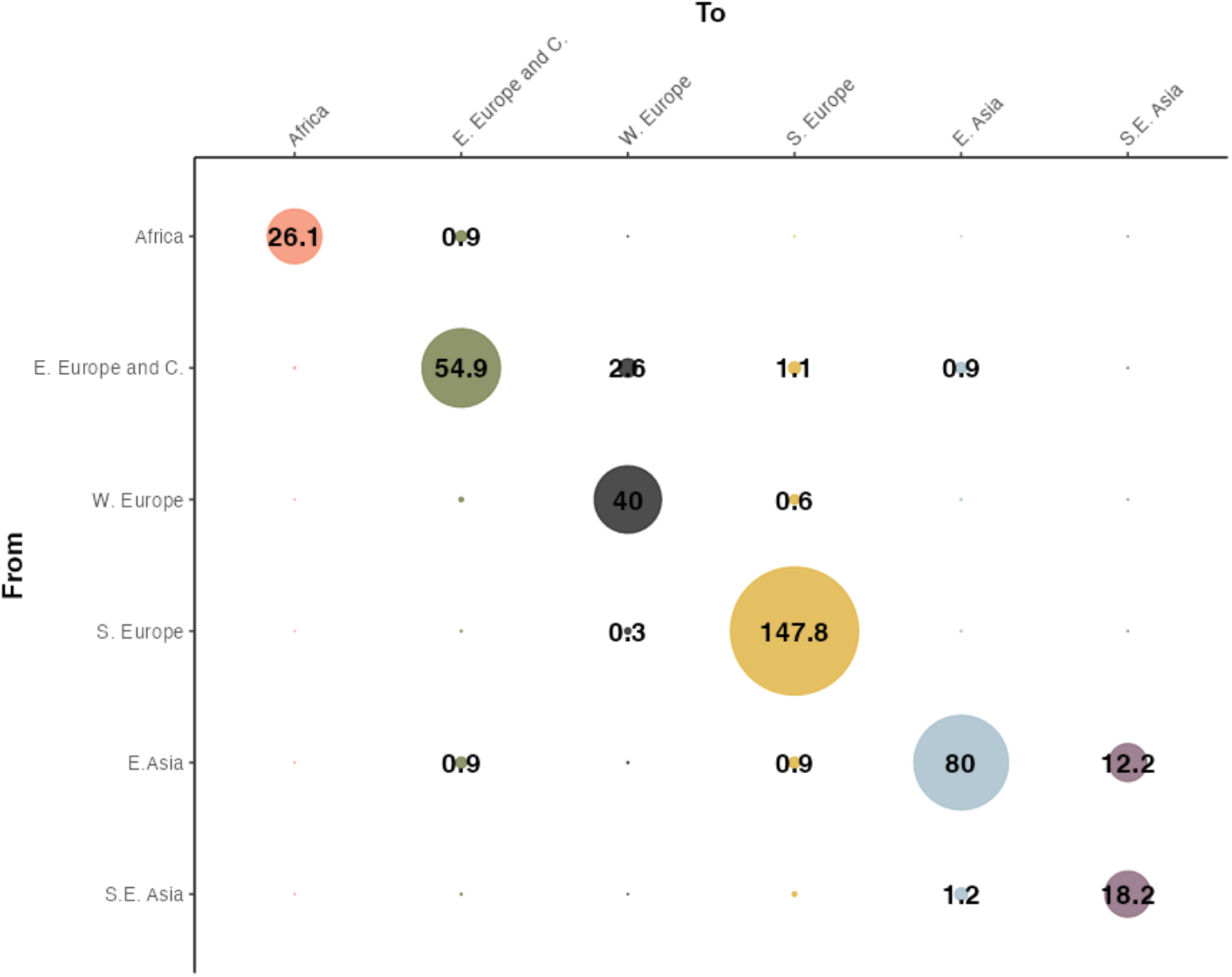
The estimated number of ASFV transitions within and between the six regions, across the 9,001 sampled phylogenetic trees. The within-region number of transitions is positively correlated with the number of whole-genome sequences available for the region (Spearman ρ = 0.94).

**Figure 4.**
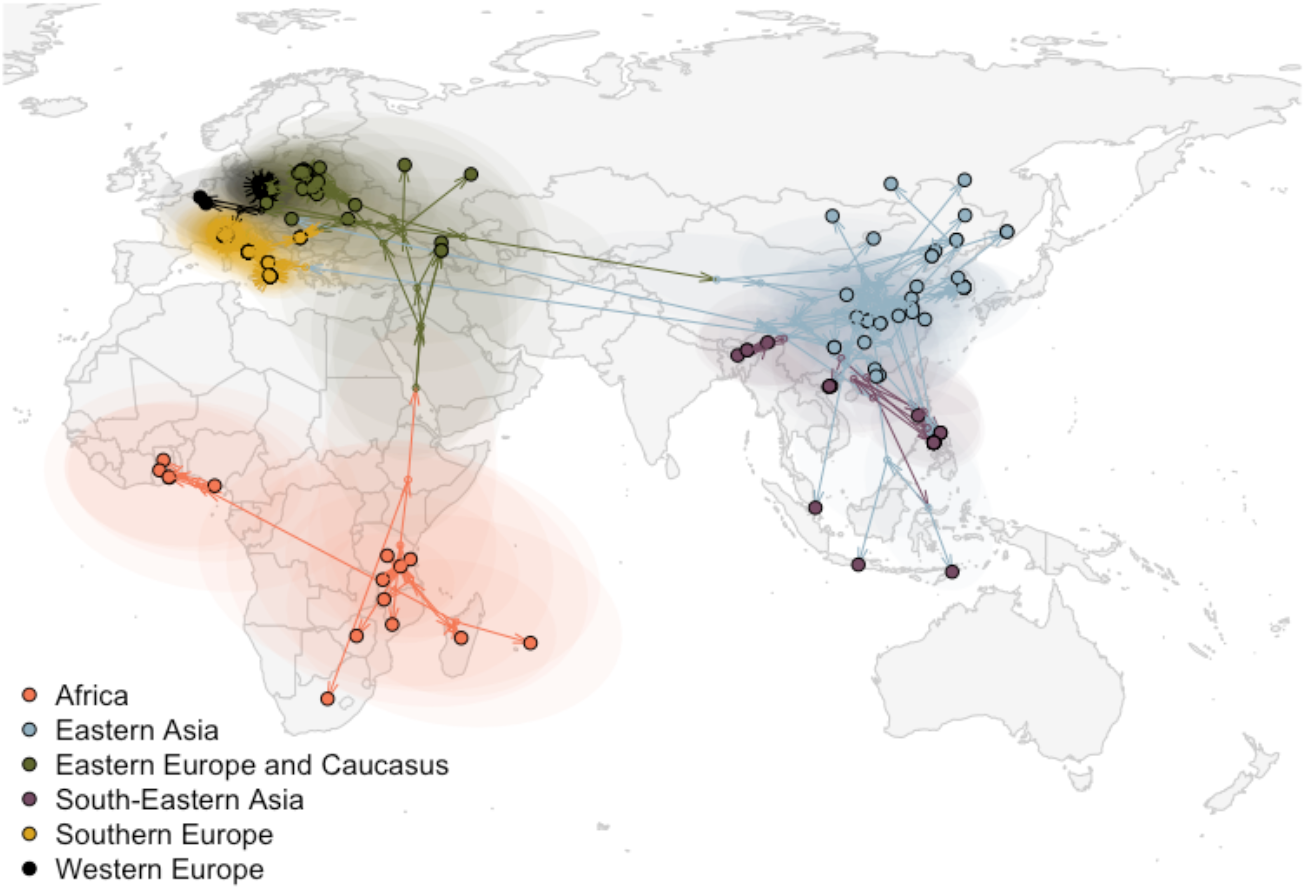
The spatially explicit MCC tree showing the global spread of ASFV estimated by BEAST. Black-circled dots represent sequences’ (i.e. tips) location (reported or centred within the smallest available administrative area), small dots represent the estimated internal nodes, arrows represent the tree’s branches, and transparent coloured areas correspond to the 80% high-posterior density. Colours correspond to the estimated (for branches or internal nodes) or observed (for tips) region.

After rewiring the branches as described, the median [95^th^ HPD] branch distance was 383.3 km [33.2 – 5,276.3], while the mode was 110km. If we consider the branches with duration less than a year (68.5% of the total), the median [95^th^ HPD] distance was 158.2[19.7 – 1,249.6] km. For branches duration up to 137, 102 and 15 days (46.7%, 40.6% and 11.2% of the total), the distances were, respectively, 146.1[18.8 – 941.9], 140.8[18.3 – 867.9], and 94.6[12.7 – 341.8] km. The cumulative distance distribution is reported in Figure 5.

**Figure 5.**
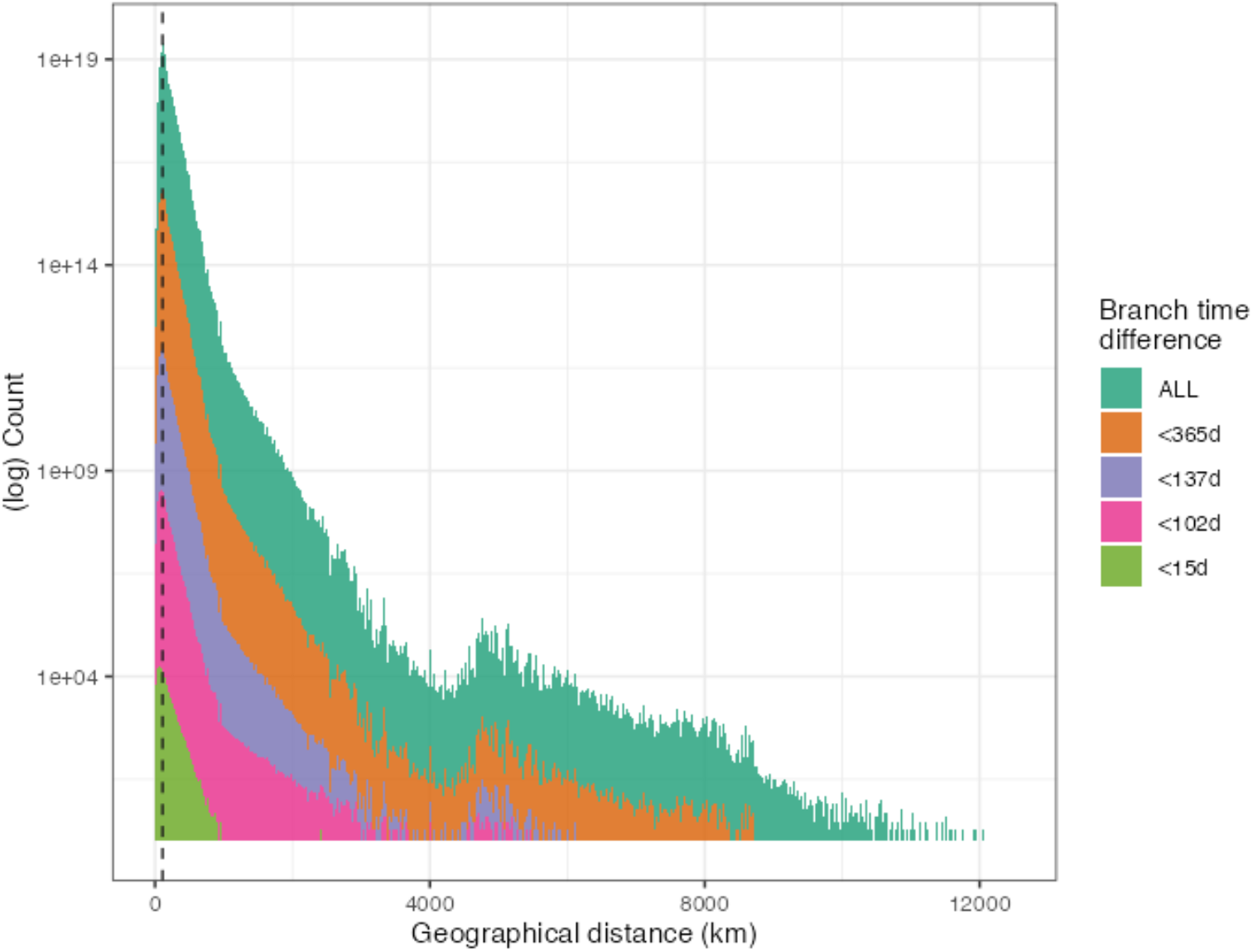
Distribution of the branches geographical distance across all 9,001 phylogenetic trees obtained in *BEAST*. The branches were sequentially rewired until both ends were located in areas where ASFV has been observed. The dashed vertical line shows the mode (110km), while colours represent the cumulative distribution of branches below a certain time threshold (teal, all time distances; orange, below 365 days; purple, below 137 days; pink, below 102 days; green, below 15 days).

### The clusters in Italy

The estimated median MRCAs of the four Italian clusters were January 2021 [HPD February 2020 – September 2021] for the North-West, July 2021 [HPD August 2020 – March 2022] for Rome municipality, January 2023 [HPD November 2022 – March 2023] for Calabria, and February 2023 [HPD November 2022 – May 2023] for Campania. Only two of the clusters were genetically similar, as in the MCC tree the Calabria clade branches off the Rome one, which in turn is associated with sequences sampled in Serbia. Conversely, the North-West cluster sequences form a clade on their own, and the sequences in the same clade are two from Belgium and one from Moldova. Finally, the Campania cluster is genetically distinct, as these strains showed similarities with the Eastern Asian clade (Figure 6).

**Figure 6.**
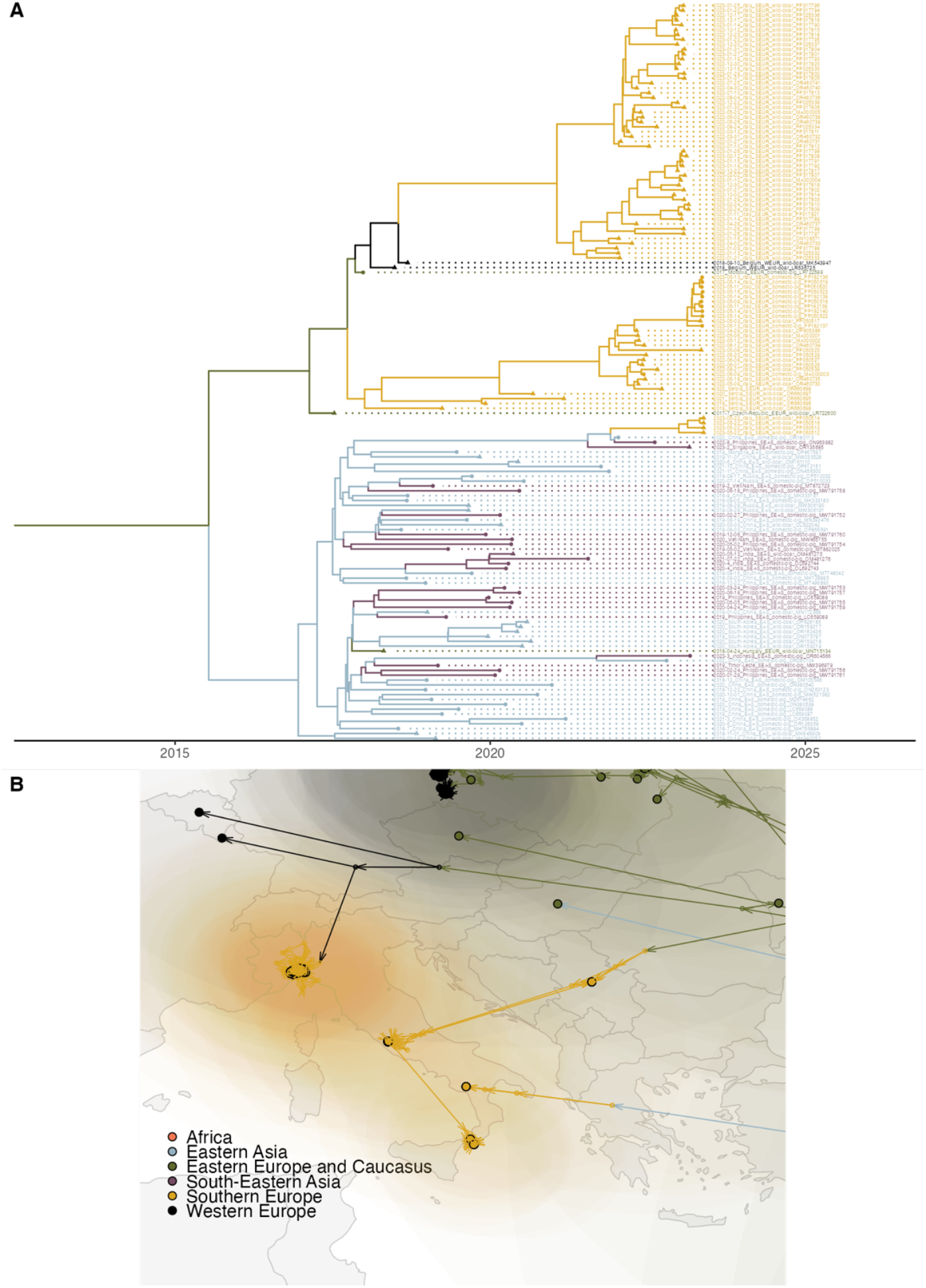
Panel A, a snapshot of the time scaled ASFV MCC tree estimated by beast, showing the two clades with the presence of WGSs sampled in Italy (European clade, top, and wider East Asia clade, bottom). Panel B, the spatially explicit MCC tree showing the spread of ASFV focused on Italy.

To confirm this result, we assessed the similarity across three individual genes, extracted from the alignment. While the B602L and I73 genes were mostly conserved, in the gene MGF 360-10L one SNP (position 986, A-to-G) was common to 67 sequences: 62 sampled in Eastern and South-Eastern Asia, one sequence from Hungary (MN715134) and four sequences from Campania, Italy (PP050512, PP050513, PP050514, and PP050516) (Figure 7). Within this clade, six sequences from South Korea (ON075797, OR159217, OR159218, OR159219, OR162436, and OR281183) and one from China (OR2910104) had another SNP. Other separate and independent clades had one SNP (*n=9* from Germany, *n=3* from Russia, MW306192, Lithuania, MK628478, and Poland, MH681419), while one sequence from Mauritius (OP781308) had two SNPs, not observed in other sequences.

**Figure 7.**
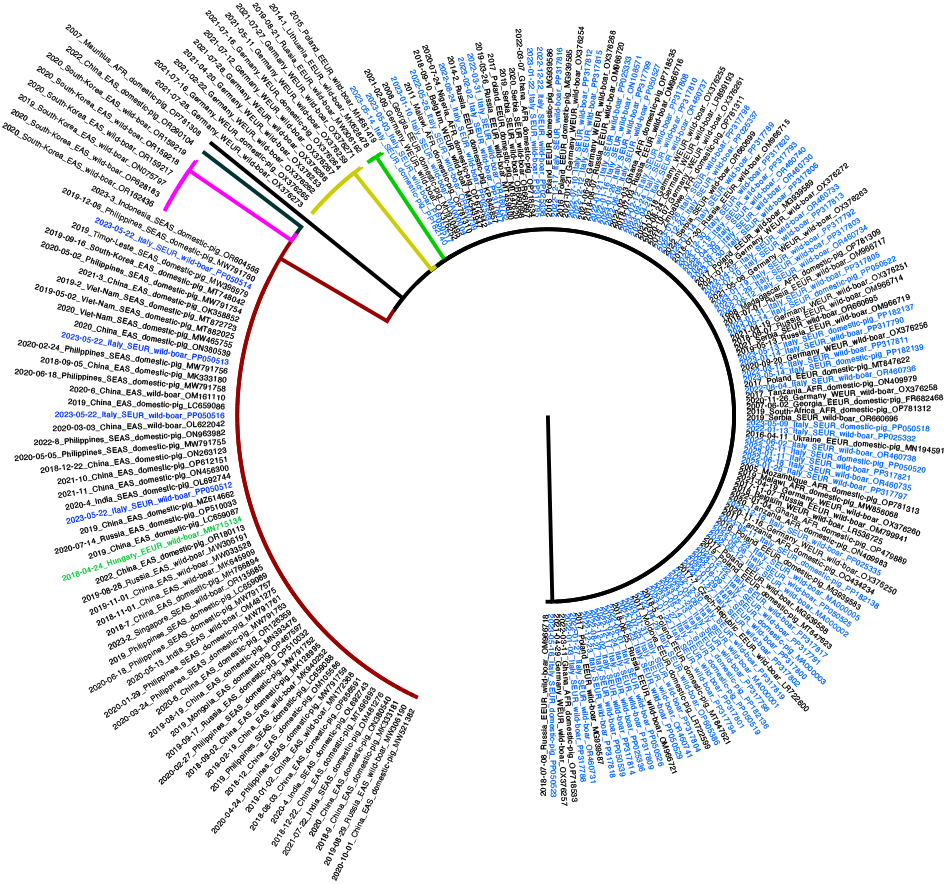
The phylogenetic tree of the ASFV MGF360-10L gene. Branch colours represent clade: main clade, black; Eastern Europe clade (*n* = 3, 1 SNP), green; German clade, gold (n = 9, 1 SNP); wide Eastern Asia clade, red (*n* = 67, 1 SNP), South Korea clade, pink (*n* = 6, 2 SNP). Single divergent sequences are coloured in black (China and Mauritius, both 2 SNPs). Italian sequences’ labels are coloured in blue, the Hungarian sequence in green (figure created with *FigTree v1*.*4*.*4*).

## Discussion

Phylodynamic approaches have been highly successful in describing the dynamics of viral infectious disease across spatial scales in human and animal populations [41–43]. Here we adopted this approach to enhance our understanding of the global spread of African Swine Fever virus, focusing on genotype II. Our findings suggest that the current ASFV panzootic is the result of long-distance anthropogenic transmissions coupled with local spread caused by either wildlife or human activities in areas where the virus encroached. A similar result was obtained by Gámbaro and coauthors [44], using a phylogeographic approach they showed how the slow ASFV dispersal dynamic was coupled with long-distance “jumps”.

The maximum clade credibility tree allowed us to pinpoint several long-distance transmission events that characterised the global spread. While the first out-of-Africa movement of genotype II, which started this panzootic, and the movement from Europe to China were already well documented [45–47], our model provided insights on other potential spreading events at the continental or intercontinental scale.

Italy is among the recently hit countries, where four clusters have been detected on the mainland since January 2022. Our analyses showed that at least three of these could likely be the results of independent introductions into the country. The clusters in the North-West and in Rome municipality were caused by distinct ASFV strains, and their most recent common ancestor dates to long before the first Italian case was observed; therefore a common origin within the country is unlikely. Previous studies employing a multi-gene approach reported similar conclusions, with the clusters in the North-West and Rome belonging to different genetic groups [22]. The sequences sampled in Rome municipality showed similarities with those from Serbia, suggesting a potential epidemiological link between the two areas: these results are also consistent with another study [44]. Conversely, the North-Western cluster isolates grouped in the same clade as one sequence from Moldova and two from Belgium. The last observed case of the Belgium outbreak was reported in mid-2019 [48], therefore a direct link between the two clusters can be excluded. Conversely, because of the absence of recent WGSs from Moldova, direct links cannot be ruled out. The MCC tree showed the Calabrian clade as a descendant of the Rome one, although the sequences in Calabria show distinctive deletions in their genome [49]. These deletions could have emerged in one of the two areas (Rome municipality or Calabria) or alternatively originated in the area that was the source of both clusters. Further investigations are required to test these hypotheses.

Surprisingly, we found the Campanian sequences were more similar to the wider Eastern Asian clade at the whole genome level than the other Italian clades, and this was also confirmed using the single gene MGF 360-10L. Specifically, we found the same SNP substitution in all East Asian sequences (both from East Asian and South-East Asia), in the Campania clade and in one sequence from Hungary, isolated in 2018. One potential explanation is that the substitution emerged in Europe, and then this variant strain was introduced to Asia and Hungary. Independently from where this strain emerged, the Campania cluster likely represents a further introduction in continental Italy. Zhang and coauthors [50] also found the Hungarian sequences branching off from the East Asian clade.

The Italian clusters need to be considered in the wider context of the ASFV circulation, specifically within the multiple virus emergence in Central, Western and Southern Europe, where many cases/outbreaks are in areas without a neighbouring affected wild boar population. Our model estimated up to seven clusters likely caused by new introductions: other than the three in Italy, Czech Republic (detected in 2017), Hungary (2018), Belgium (2018), Western Poland (2019). The latter further expanded to Germany, where it was first detected in 2020 [51]. We are also aware of at least two further introductions not reported in our study, in Sweden [52] and Baden-Wurttemberg (Germany, 2022, personal communication). This situation is mirrored in South-Eastern Asia, where the pathogen was introduced in several countries, including on many islands, in the span of a few years. The circulation in this area is well represented in the transition matrix, which shows how the transitions between East and South-East Asia were similar to the within-region ones. Another notable long-distance movement of ASFV was from South-Eastern to Western Africa, where it emerged in Nigeria in 2020 [53,54] and Ghana in 2022 [55], although according to our estimates, it might have been circulating in the region as early as 2017.

The current analysis presented many challenges related to the pathogen itself and to its epidemiology. While phylogenetic and phylogeographic approaches have been mostly used to tackle fast mutating RNA viruses, such as avian flu[56] and Porcine Reproductive and Respiratory Syndrome virus[57], recent developments showed these can be applied to slow-evolving pathogens, such as *Staphylococcus aureus[58]* or *Mycobacterium bovis*[59,60]. ASFV could be considered in the middle of these variation-wise, with a substitution rate of approximately two substitutions/genome/year. The high similarity of many sequences, however, could have caused a low posterior probability value in some branches of the MCC tree (Figure S4): this was the case for smaller clusters, like the North-Western Italian clade, but also for the wider Eastern Asian clade. To avoid further biases in the MCC tree, discrete and continuous traits were added in a second phase.

Following the exclusion of internal nodes in presumed ASFV-free regions, and subsequent branch rewiring, we analysed the resulting branch distance distribution across the 9,001 trees. Branches with durations equal or lower than 137 days, which could potentially allow the virus survival in cured or refrigerated pork, exceeded 900 km (Figure 5), further highlighting the potential role of contaminated material movement in ASFV spread. If we consider survival times as short as 15 days, also comparable to the virus generation time, the median distance was approximately 95 km, and branches could be as long as 340 km, far beyond the potential reach of infected wildlife movements.

Similarly to two previous studies [50,61] we estimated a clock rate in the order of 1-1.5×10^-5^ (1.07×10^-5^, compared to 1.14×10^-5^ and 1.31×10^-5^), but higher than another recent study[44] (0.57×10^-5^). We employed an exponential relaxed clock model, whereas the first two studies used a strict model and the latter a logistic distributed clock. These results are reflected in the Eurasian epidemic MRCA estimate: our result (November 2005 [HPD September 2003 – May 2007]) was similar to Zhang and co-authors[50], but with a narrower HPD interval (August 2005 [HPD April 2000–January 2007), but wider than Gámbaro and co-authors[44] (HPD August 2004–September 2007).

A limitation is the notable absence of WGSs from Eastern Europe between 2008 and 2014, and the relatively small number of sequences from countries infected earlier in the epidemic. Isolates from these areas, and eventually time periods, could provide further insights on the early spread of the disease and therefore improve the robustness of the MCC tree estimates. Another important question is whether the introduction of ASFV into a naïve population is leading to the emergence of new strains or variants [40], in particular in the context where ticks are not involved in the spread and maintenance of the disease.

While our findings provide further insights on ASFV genotype II global circulation, a potential extension of this work could help to uncover social, economic or environmental drivers - beyond the presence of wild boars-that enable its introduction and establishment in new regions. Determining which of these factors are associated with long distance transmissions could be crucial to improve the biosecurity and surveillance in those areas deemed at higher risk.

## Conclusions

In conclusion, our phylogenetic model of ASFV genotype II reveals a complex dynamic including importations layered over local transmissions in both wild and domestic swine. The uncovering of multiple introductions into Europe -including at least three into Italy- and the potential genetic links to East Asian lineages expose critical blind spots. To bridge these gaps, genomic surveillance of the virus needs to be expanded and improved across under-sampled regions, and to consider these data with ecological, trade and socio-economic information, using complementary epidemiological approaches. Only through such an integrated, high-resolution monitoring framework can we sharpen real-time risk assessments, design context-specific biosecurity measures, and ultimately stop ASFV’s expansion.

## Acknowledgements

The authors thank Cesare Cammà and Maurilia Marcacci from the Istituto Zooprofilattico Sperimentale dell’Abruzzo e del Molise G. Caporale (Teramo, Italy) for the assistance and the whole-genome sequencing of the Italian isolates. For the purpose of open access, the author has applied a CC-BY public copyright licence to any Author Accepted Manuscript version arising from this submission.

## Data availability

All the African Swine Fever virus genomes used in this study are deposited in NCBI GenBank (see Supplementary material for accession numbers).

## References

[1] S. You, T. Liu, M. Zhang, X. Zhao, Y. Dong, B. Wu et al., African swine fever outbreaks in China led to gross domestic product and economic losses, Nat Food 2 (2021), pp. 802–808.

[2] J.K. Niemi, Impacts of African Swine Fever on Pigmeat Markets in Europe, Frontiers in Veterinary Science 7 (2020), pp. 1–11.

[3] E. Meijaard, A. Erman, M. Ancrenaz and B. Goossens, Pig virus imperils food security in Borneo, Science 383 (2024), pp. 267–267.

[4] R. Eustace Montgomery, On A Form of Swine Fever Occurring in British East Africa (Kenya Colony), Journal of Comparative Pathology and Therapeutics 34 (1921), pp. 159–191.

[5] L.K. Dixon, H. Sun and H. Roberts, African swine fever, Antiviral Research 165 (2019), pp. 34–41.

[6] H. Zhang, S. Zhao, H. Zhang, Z. Qin, H. Shan and X. Cai, Vaccines for African swine fever: an update, Front. Microbiol. 14 (2023).

[7] S. Blome, K. Franzke and M. Beer, African swine fever – A review of current knowledge, Virus Research 287 (2020), pp. 198099.

[8] J.M. Sánchez-Vizcaíno, L. Mur, J.C. Gomez-Villamandos and L. Carrasco, An Update on the Epidemiology and Pathology of African Swine Fever, Journal of Comparative Pathology 152 (2015), pp. 9–21.

[9] S. Petrini, F. Feliziani, C. Casciari, M. Giammarioli, C. Torresi and G.M. De Mia, Survival of African swine fever virus (ASFV) in various traditional Italian dry-cured meat products, Preventive Veterinary Medicine 162 (2019), pp. 126–130.

[10] S. Costard, B. Wieland, W. de Glanville, F. Jori, R. Rowlands, W. Vosloo et al., African swine fever: how can global spread be prevented?, Philosophical Transactions of the Royal Society B: Biological Sciences 364 (2009), pp. 2683–2696.

[11] World Animal Health Information System - WOAH (formerly OIE), WOAH - World Organisation for Animal Health.

[12] X. Zhou, N. Li, Y. Luo, Y. Liu, F. Miao, T. Chen et al., Emergence of African Swine Fever in China, 2018, Transboundary and Emerging Diseases 65 (2018), pp. 1482– 1484.

[13] E. Mighell and M.P. Ward, African Swine Fever spread across Asia, 2018–2019, Transboundary and Emerging Diseases 68 (2021), pp. 2722–2732.

[14] Q. Shao, R. Li, Y. Han, D. Han and J. Qiu, Temporal and Spatial Evolution of the African Swine Fever Epidemic in Vietnam, International Journal of Environmental Research and Public Health 19 (2022), pp. 8001.

[15] R.P. Jean-Pierre, A.D. Hagerman and K.M. Rich, An analysis of African Swine Fever consequences on rural economies and smallholder swine producers in Haiti, Front. Vet. Sci. 9 (2022).

[16] V.R. Brown, R.S. Miller, K.M. Pepin, K.M. Carlisle, M.A. Cook, C.F. Vanicek et al., African swine fever at the wildlife-livestock interface: challenges for management and outbreak response within invasive wild pigs in the United States, Front. Vet. Sci. 11 (2024).

[17] N. Mazur-Panasiuk, J. Żmudzki and G. Woźniakowski, African swine fever virus – persistence in different environmental conditions and the possibility of its indirect transmission, Journal of Veterinary Research 63 (2019), pp. 303–310.

[18] L. Mur, B. Martínez-López and J.M. Sánchez-Vizcaíno, Risk of African swine fever introduction into the European Union through transport-associated routes: returning trucks and waste from international ships and planes, BMC Vet Res 8 (2012), pp. 149.

[19] C. Iscaro, A. Dondo, L. Ruocco, L. Masoero, M. Giammarioli, S. Zoppi et al., January 2022: Index case of new African Swine Fever incursion in mainland Italy, Transboundary and Emerging Diseases 69 (2022), pp. 1707–1711.

[20] M. Giammarioli, D. Alessandro, C. Cammà, L. Masoero, C. Torresi, M. Marcacci et al., Molecular Characterization of the First African Swine Fever Virus Genotype II Strains Identified from Mainland Italy, 2022, Pathogens 12 (2023), pp. 372.

[21] M. Giammarioli, M. Marcacci, M.T. Scicluna, A. Cersini, C. Torresi, V. Curini et al., Complete Genome of African Swine Fever Virus Genotype II in Central Italy, Microbiology Resource Announcements (2023).

[22] M. Giammarioli, C. Torresi, R. Biccheri, C. Cammà, M. Marcacci, A. Dondo et al., Genetic Characterization of African Swine Fever Italian Clusters in the 2022-2023 Epidemic Wave by a Multi-Gene Approach, Viruses 16 (2024), pp. 1185.

[23] National Center for Biotechnology Information. Available at https://www.ncbi.nlm.nih.gov/.

[24] K. Katoh and D.M. Standley, MAFFT Multiple Sequence Alignment Software Version 7: Improvements in Performance and Usability, Mol Biol Evol 30 (2013), pp. 772–780.

[25] M.A. Suchard, P. Lemey, G. Baele, D.L. Ayres, A.J. Drummond and A. Rambaut, Bayesian phylogenetic and phylodynamic data integration using BEAST 1.10, Virus Evolution 4 (2018), pp. 1–5.

[26] FigTree. Available at http://tree.bio.ed.ac.uk/software/Figtree/.

[27] R Core Team, R: A Language and Environment for Statistical Computing, R Foundation for Statistical Computing, Vienna, Austria, 2024.

[28] Empres-Plus. Available at https://empres-i.apps.fao.org/general.

[29] A.M.M. Stoian, J. Zimmerman, J. Ji, T.J. Hefley, S. Dee, D.G. Diel et al., Half-Life of African Swine Fever Virus in Shipped Feed - Volume 25, Number 12—December 2019 - Emerging Infectious Diseases journal - CDC.

[30] D.A.G. Chapman, A.C. Darby, M.D. Silva, C. Upton, A.D. Radford and L.K. Dixon, Genomic Analysis of Highly Virulent Georgia 2007/1 Isolate of African Swine Fever Virus - Volume 17, Number 4—April 2011 - Emerging Infectious Diseases journal - CDC.

[31] C. Gallardo, N. Casado, A. Soler, I. Djadjovski, L. Krivko, E. Madueño et al., A multi gene-approach genotyping method identifies 24 genetic clusters within the genotype II-European African swine fever viruses circulating from 2007 to 2022, Front. Vet. Sci. 10 (2023).

[32] A. Mazloum, A. van Schalkwyk, R. Chernyshev, A. Igolkin, L. Heath and A. Sprygin, A Guide to Molecular Characterization of Genotype II African Swine Fever Virus: Essential and Alternative Genome Markers, Microorganisms 11 (2023), pp. 642.

[33] C. Gallardo, D.M. Mwaengo, J.M. Macharia, M. Arias, E.A. Taracha, A. Soler et al., Enhanced discrimination of African swine fever virus isolates through nucleotide sequencing of the p54, p72, and pB602L (CVR) genes, Virus Genes 38 (2009), pp. 85– 95.

[34] Y. Liu, Z. Shen, Z. Xie, Y. Song, Y. Li, R. Liang et al., African swine fever virus I73R is a critical virulence-related gene: A potential target for attenuation, Proceedings of the National Academy of Sciences 120 (2023), pp. e2210808120.

[35] T.G. Burrage, Z. Lu, J.G. Neilan, D.L. Rock and L. Zsak, African Swine Fever Virus Multigene Family 360 Genes Affect Virus Replication and Generalization of Infection in Ornithodoros porcinus Ticks, Journal of Virology 78 (2004), pp. 2445–2453.

[36] D. Li, J. Peng, J. Wu, J. Yi, P. Wu, X. Qi et al., African swine fever virus MGF-360-10L is a novel and crucial virulence factor that mediates ubiquitination and degradation of JAK1 by recruiting the E3 ubiquitin ligase HERC5, mBio 14 (2023), pp. e00606–23.

[37] S.F. Altschul, W. Gish, W. Miller, E.W. Myers and D.J. Lipman, Basic local alignment search tool, Journal of molecular biology 215 (1990), pp. 403–410.

[38] B.Q. Minh, H.A. Schmidt, O. Chernomor, D. Schrempf, M.D. Woodhams, A. Von Haeseler et al., IQ-TREE 2: New Models and Efficient Methods for Phylogenetic Inference in the Genomic Era, Molecular Biology and Evolution 37 (2020), pp. 1530– 1534.

[39] M.S. Gill, P. Lemey, N.R. Faria, A. Rambaut, B. Shapiro and M.A. Suchard, Improving bayesian population dynamics inference: A coalescent-based model for multiple loci, Molecular Biology and Evolution 30 (2013), pp. 713–724.

[40] J.H. Forth, S. Calvelage, M. Fischer, J. Hellert, J. Sehl-Ewert, H. Roszyk et al., African swine fever virus – variants on the rise, Emerging Microbes & Infections 12 (2023), pp. 2146537.

[41] C. Guinat, T. Vergne, A. Kocher, D. Chakraborty, M.C. Paul, M. Ducatez et al., Ecology & Evolution What can phylodynamics bring to animal health research ?, Trends in Ecology & Evolution (2021), pp. 1–11.

[42] P. Lemey, A. Rambaut, T. Bedford, N. Faria, F. Bielejec, G. Baele et al., Unifying Viral Genetics and Human Transportation Data to Predict the Global Transmission Dynamics of Human Influenza H3N2, PLoS Pathogens 10 (2014), .

[43] F. Duchatel, B.M. de C. Bronsvoort and S. Lycett, Phylogeographic Analysis and Identification of Factors Impacting the Diffusion of Foot-and-Mouth Disease Virus in Africa, Front. Ecol. Evol. 7 (2019).

[44] F. Gámbaro, L.C. Goatley, T.J. Foster, C. Tennakoon, G.L. Freimanis, S. Van Borm et al., Exploiting Viral DNA Genomes to Explore the Dispersal History of African Swine Fever Genotype II Lineages in Europe, Genome Biology and Evolution 17 (2025), pp. evaf102.

[45] N.N. Gaudreault, D.W. Madden, W.C. Wilson, J.D. Trujillo and J.A. Richt, African Swine Fever Virus: An Emerging DNA Arbovirus, Front. Vet. Sci. 7 (2020).

[46] E.P. Njau, J.-B. Domelevo Entfellner, E.M. Machuka, E.N. Bochere, S. Cleaveland, G.M. Shirima et al., The first genotype II African swine fever virus isolated in Africa provides insight into the current Eurasian pandemic, Sci Rep 11 (2021), pp. 13081.

[47] R.J. Rowlands, V. Michaud, L. Heath, G. Hutchings, C. Oura, W. Vosloo et al., African Swine Fever Virus Isolate, Georgia, 2007 - Volume 14, Number 12—December 2008 - Emerging Infectious Diseases journal - CDC.

[48] S. Dellicour, D. Desmecht, J. Paternostre, C. Malengreaux, A. Licoppe, M. Gilbert et al., Unravelling the dispersal dynamics and ecological drivers of the African swine fever outbreak in Belgium, Journal of Applied Ecology 57 (2020), pp. 1619–1629.

[49] C. Torresi, R. Biccheri, C. Cammà, C. Gallardo, M. Marcacci, S. Zoppi et al., Genome-Wide Approach Identifies Natural Large-Fragment Deletion in ASFV Strains Circulating in Italy During 2023, Pathogens 14 (2025), pp. 51.

[50] Y. Zhang, Q. Wang, Z. Zhu, S. Wang, S. Tu, Y. Zhang et al., Tracing the Origin of Genotype II African Swine Fever Virus in China by Genomic Epidemiology Analysis, Transboundary and Emerging Diseases 2023 (2023), pp. 4820809.

[51] C. Sauter-Louis, J.H. Forth, C. Probst, C. Staubach, A. Hlinak, A. Rudovsky et al., Joining the club: First detection of African swine fever in wild boar in Germany, Transboundary and Emerging Diseases 68 (2021), pp. 1744–1752.

[52] E. Chenais, V. Ahlberg, K. Andersson, F. Banihashem, L. Björk, M. Cedersmyg et al., First Outbreak of African Swine Fever in Sweden: Local Epidemiology, Surveillance, and Eradication Strategies, Transboundary and Emerging Diseases 2024 (2024), pp. 6071781.

[53] A.J. Adedeji, P.D. Luka, R.B. Atai, T.A. Olubade, D.A. Hambolu, M.A. Ogunleye et al., First-Time Presence of African Swine Fever Virus Genotype II in Nigeria, Microbiology Resource Announcements 10 (2021), pp. 10.1128/mra.00350-21.

[54] A. Ambagala, K. Goonewardene, L. Lamboo, M. Goolia, C. Erdelyan, M. Fisher et al., Characterization of a Novel African Swine Fever Virus p72 Genotype II from Nigeria, Viruses 15 (2023), pp. 915.

[55] E. Spinard, A. Rai, J. Osei-Bonsu, V. O’Donnell, P.T. Ababio, D. Tawiah-Yingar et al., The 2022 Outbreaks of African Swine Fever Virus Demonstrate the First Report of Genotype II in Ghana, Viruses 15 (2023), pp. 1722.

[56] S.J. Lycett, A. Pohlmann, C. Staubach, V. Caliendo, M. Woolhouse, M. Beer et al., Genesis and spread of multiple reassortants during the 2016/2017 H5 avian influenza epidemic in Eurasia, Proceedings of the National Academy of Sciences 117 (2020), pp. 20814–20825.

[57] D.N. Makau, I.A.D. Paploski, C.A. Corzo and K. VanderWaal, Dynamic network connectivity influences the spread of a sub-lineage of porcine reproductive and respiratory syndrome virus, Transboundary and Emerging Diseases 69 (2022), pp. 524– 537.

[58] G. Yebra, J.D. Harling-Lee, S. Lycett, F.M. Aarestrup, G. Larsen, L.M. Cavaco et al., Multiclonal human origin and global expansion of an endemic bacterial pathogen of livestock, Proceedings of the National Academy of Sciences 119 (2022), pp. e2211217119.

[59] J. Crispell, C.H. Benton, D. Balaz, N.D. Maio, A. Akhmetova, A. Allen et al., Combining genomics and epidemiology to analyse bidirectional transmission of Mycobacterium bovis in a multi-host system, eLife 8 (2019), pp. 1–36.

[60] G. Rossi, B.B.-J. Shih, N.F. Egbe, P. Motta, F. Duchatel, R.F. Kelly et al., Unraveling the epidemiology of Mycobacterium bovis using whole-genome sequencing combined with environmental and demographic data, Frontiers in Veterinary Science 10 (2023).

[61] Z.-J. Shen, H. Jia, C.-D. Xie, J. Shagainar, Z. Feng, X. Zhang et al., Bayesian Phylodynamic Analysis Reveals the Dispersal Patterns of African Swine Fever Virus, Viruses 14 (2022), pp. 889.

