## Supplementary material for "A phylogenetic contribution to understanding the panzootic spread of African swine fever: from the global to the local scale"

***Metadata preparation***

To run the phylodynamic analyses of the African Swine Fever virus (ASFV), metadata associated with the whole-genome sequences was required. In particular, we used: sample date, geographical location and host (wild boar or domestic pig). All the metadata of the 82 Italian sequences was available, collected by the local veterinarian health authorities together with the infected animals tissue. Only part of the 146 sequences retrieved from online repositories reported the available metadata, and they were obtained using the entrez*_summary()* function of the *rentrex R* package [1]. For the others, the assigned coordinates were the centroids of the smallest administrative area associated with each sequence. Four WGSs available from Japan were obtained from contaminated meat seized at two airports [2]. In this case, we assigned the coordinates to the city of each flight origins, two from China (Shangai and Qingdao) and two from the Philippines (both Manila). The *BEAST* software, that we used for the main analyses, can handles partial dates (i.e. only year or year and month) by assigning an uncertainty interval.

We also assigned a region to each sequence according to the United Nations Geoscheme sub-regions[3]. However, to reach enough sequences per region for the analysis to be robust (15), some of them were aggregated, specifically: African sequences were aggregated in one region (*n*=15); Transcaucasian (*n*=3, two from Georgia and one from Armenia) and Northern Europe (*n*=2, from Estonia and Lithuania) sequences were assigned to Eastern Europe and Caucasus; South Asia (*n*=4, from India) were assigned to South-Eastern Asia; finally Eastern Russia sequences (*n*=4) were assigned to Eastern Asia, as they were sampled in the far east part of the country. The sequences (observed or estimated) locations are reported in Fig. 1, while a timeline of the sequences sampling is shown in Figure S1.

***Preliminary analyses and model selection***

Before running the models, the 228 African Swine Fever virus whole genome sequences (WGS) went through a preliminary quality control stage. We run the software *IQ-TREE2*[4] to test for the best substitution model, create a maximum-likelihood tree and identify potential duplicates and poor quality sequences.

Two sequences (LR881473 from Armenia, LS478113 from Estonia) were removed as they returned poor results overall. The untrimmed alignment length was 198,919 bp, while the trimmed one was 188,125 bp. After a further manual check, it was reduced to 184,545 bp (most of the removals were empty columns following the removal of LR881473 and LS478113 from the alignment).

The software *IQ-TREE2* outputs showed that ON400500 (China) was not viable for analysis, as it did not pass the composition test. Furthermore, seven pairs of sequences were found identical. While three pairs (two from Italy and one from China) were sampled on separate days and locations, the same could not be confirmed for other four pairs, thus one sequence for each pair was removed (PP050528 and PP050521 from Italy, MN393477 from China, and NC_044948 from Russia) to avoid potential duplicates. The analysis of the root-to-tip distance also highlighted two further outliers' sequences, which were not considered for the following analyses: OQ737679, OM105587, and MW656282 from China, and MG939584 from Poland. The latter two sequences were excluded also from another study [5]. The final alignment we used for the analyses included 217 WGSs.

To test whether the sampled pathogen population had enough genetic variability to run the phylogenetic analyses, we assessed the genetic signal using the maximum-likelihood tree resulted from *IQ-TREE2* with the software *Tempest v1.5.3* [6]. Specifically, we tested whether a lineal model of root-to-tip distance vs. the temporal distance had a positive slope. For the Maximum Likelihood tree computed by *IQ-TREE2* using the final 217-WGS alignment, we obtained a positive correlation between the root-to-tip distance and the temporal distance (slope $0.52\times10^{-5}$ and $R^2$ 0.20).

Finally, the software *OpenRDP[7]* was used to search for recombination signals. None of the sequences was identified as recombinant by more than two of the algorithms run by the software, therefore we considered this result inconclusive.

We tested many preliminary *BEAST* runs [8] to select the best different clock rate models (strict, relaxed exponential, relaxed logistic), clock rate prior distributions, and different population size models, or tree priors (constant population, exponential growth, and Bayesian SkyGrid [9]). In the preliminary models, the chain length of $10^8$ steps, sampled every $10^4$ steps were used to obtain 1,001 trees. We evaluated whether the models converged, using the Effective Sample Size (ESS) over 200 as main criterion, with the software *Tracer v.1.7.2* [10]. To compare multiple converged models we

used the Marginal Likelihood Estimation (MLE), calculated with the Path sample and Stepping-stone sample [11].
The best substitution model obtained by *IQ-TREE2[4]*, and according to the Bayesian information criterion, was the Hasegawa-Kishino-Yano (HKY), with invariable sites and empirical base frequency. However, our selection process showed that the addition of Gamma-distributed heterogeneity model improved the model in *BEAST* (Table S2). The best clock rate model was the exponentially distributed relaxed distribution, while the shape of the prior did not impact significantly the results. To allow flexibility in the pathogen effective population size (i.e. tree prior) we used a Bayesian SkyGrid model, with 54 estimates and 27 years as time to last transition. We tested this against other models, as well as different SkyGrid parameters (Table S2). We also run a model using parameters used in other studies found in the literature [12–14].
Once the best phylogenetic model was identified, the model was run again using alternative chain lengths and random number seeds to improve variability. We ran replicates of the model six further times and combined the results of the independent runs (i.e. seven in total: two with chain length $10^8$, three with chain length $2\times10^8$, and one of each with chain length $4\times10^8$ and $10^9$). A sample of 300 trees was collected from the posterior 10,001 trees distribution of each run and combined to obtain 2,100 trees. These posterior tree samples were used as input for a further run in *BEAST* to generate a new trees distribution with trait partitions. Two of these traits were discrete, identifying the isolates region (see Tab. 1) and host (wild or domestic), and one continuous (geographical coordinates). We set an asymmetric model for the discrete traits, so that the transitions between them were calculated independently for each direction, and a Brownian random walk model for the spatial model. The chain length was again $10^8$, sampled every $10^4$ to obtain a final 10,001 trees. Of these, the first 1,000 were discarded as burn-in, so to focus our analysis on the remaining 9,001.

***Single gene analysis result: B602L and I73R genes***

We built Maximum Likelihood trees on single genes. The B602L gene was mostly conserved, with only a clade of five sequences from Western Africa (four from Ghana, OP718533, OP718534, OP718535, and OP479889, one from Nigeria, OP672342) with two single nucleotide polymorphisms (SNPs), an additional one (Ukraine, MN194591) with two single-point deletions, another one with a single-point deletion (Russia, KP843857) and two with one SNP each (from Russia, MW306192, and China, OM105586). Similarly, the I73R was conserved across the alignment, with only one sequence reporting three SNPs (China, MK940252).

### Supplementary figures and tables

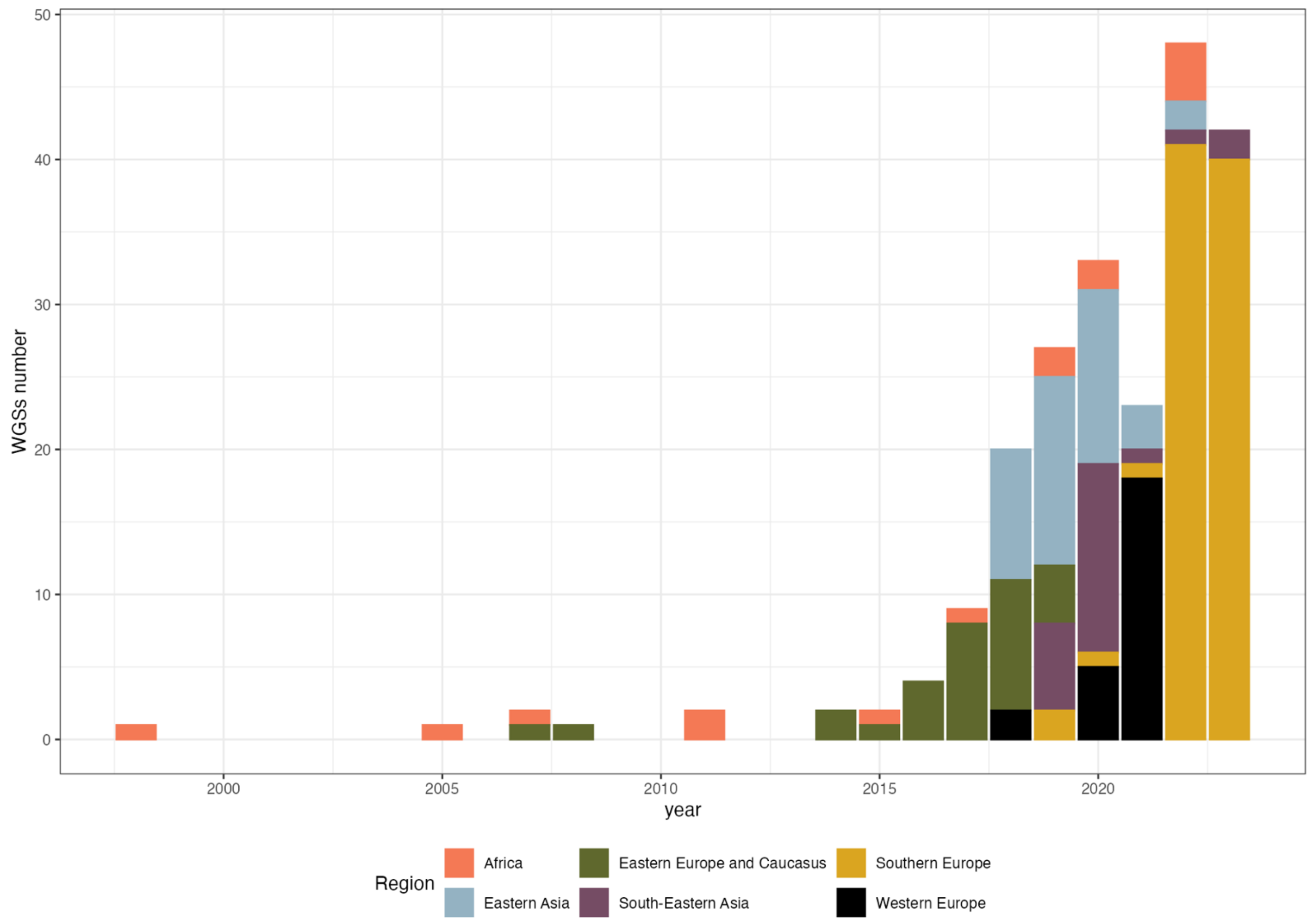


**Figure S1**. Sampling timeline of the 228 African Swine Fever virus (ASFV) whole-genome sequences considered for this study. Colours correspond to different regions (Africa, red; Eastern Asia, blue; South-Eastern Asia, purple; Eastern Europe and Caucasus, green; Southern Europe, yellow; Western Europe, black).

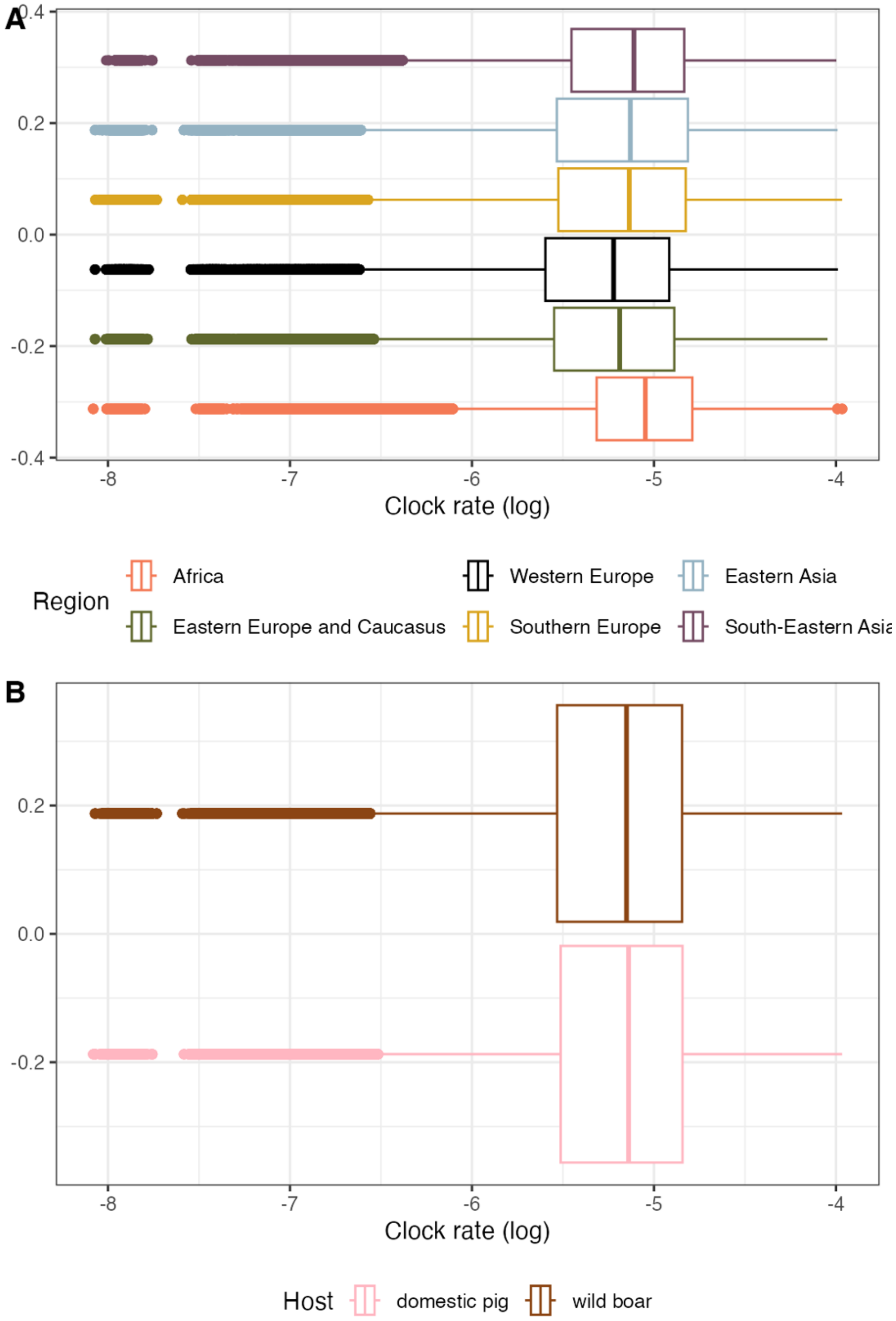


**Figure S2**. The estimated clock rate calculated for the within-region (panel A) and within-host (B) transitions. In panel A colours correspond to different regions (Africa, red; Eastern Asia, blue; South-Eastern Asia, purple; Eastern Europe and Caucasus, green; Southern Europe, yellow; Western Europe, black), while in panel B to host (pink for domestic pigs, brown for wild boar or feral pigs).

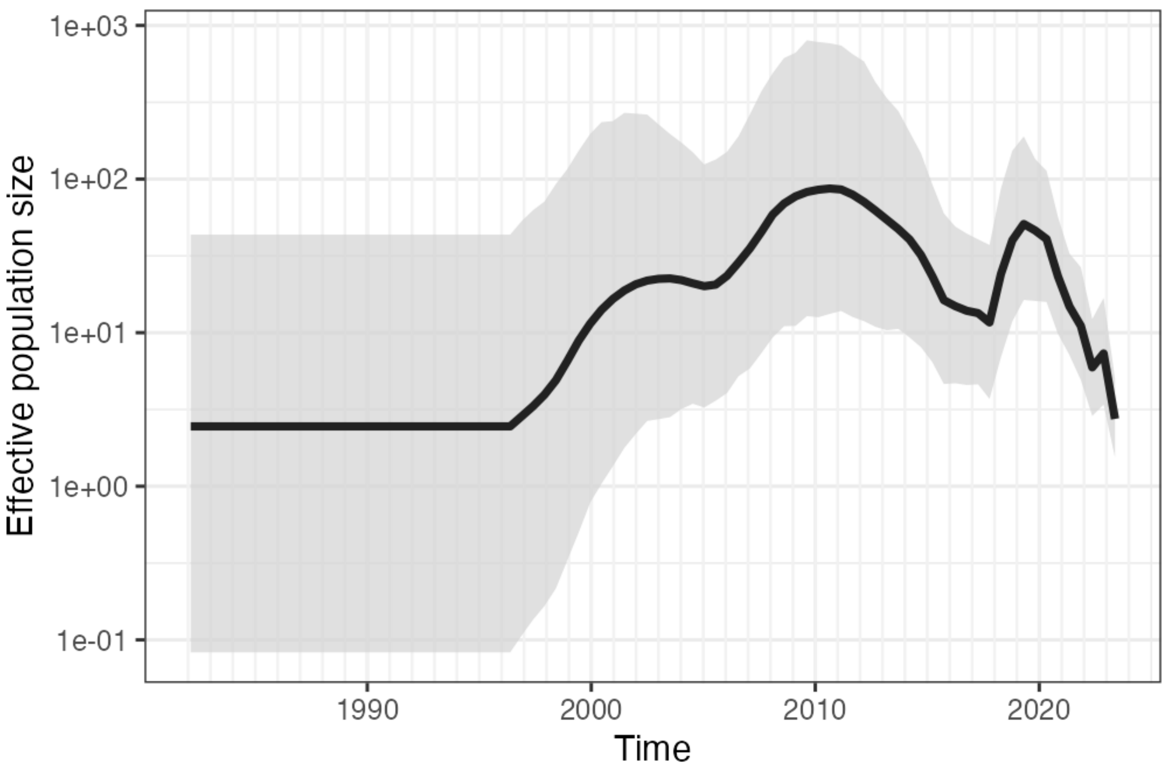


**Figure S3**. The effective population size of the ASFV estimated by BEAST. The phylogenetic tree prior model was SkyGrid with 54 estimates and 27 years as time to last transition. Starting from the late 1990s (the same period the oldest sequence was sampled), we observed an initial expansion of the effective population size until early 2000s, when the virus was likely still in Africa. We then observed two main peaks: one between 2010 and 2011, and the second between 2018 and 2019. The first peak might correspond to the initial spread in Eastern Europe, before its introduction in the European Union, while the second might be caused by the widespread outbreaks in China. The decline of the effective population size after 2021 might be caused by the similarity of the sequences available those years, despite a higher number of them in our dataset (see Fig. S1).

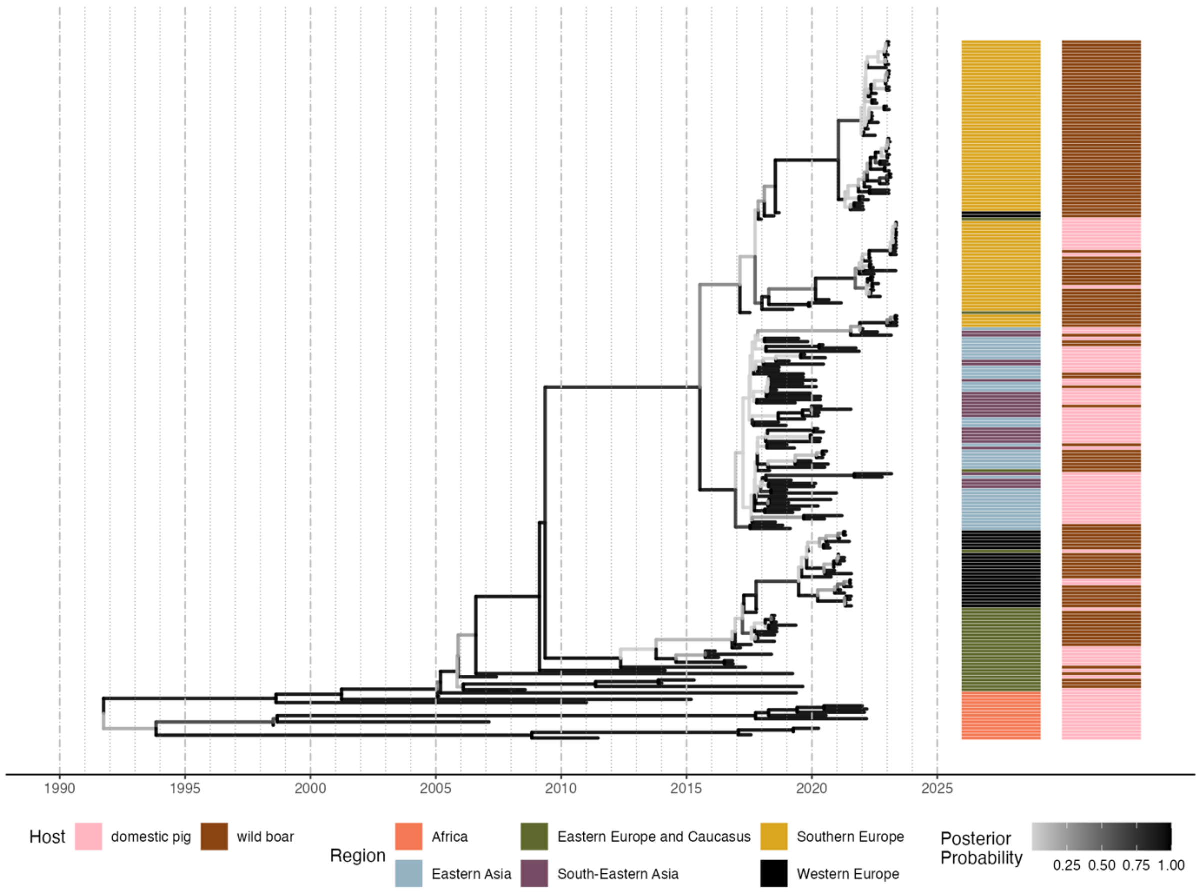


**Figure S4**. The Maximum Clade Credibility (MCC) tree of the 217 African Swine fever whole-genome sequences used in the study. Branches are coloured according to the MMC branches posterior probability calculated in BEAST. The two columns on the right report, respectively, the tips region (Africa, red; Eastern Asia, blue; South-Eastern Asia, purple; Eastern Europe and Caucasus, green; Southern Europe, yellow; Western Europe, black), and host (pink for domestic pigs, brown for wild boar or feral pigs).

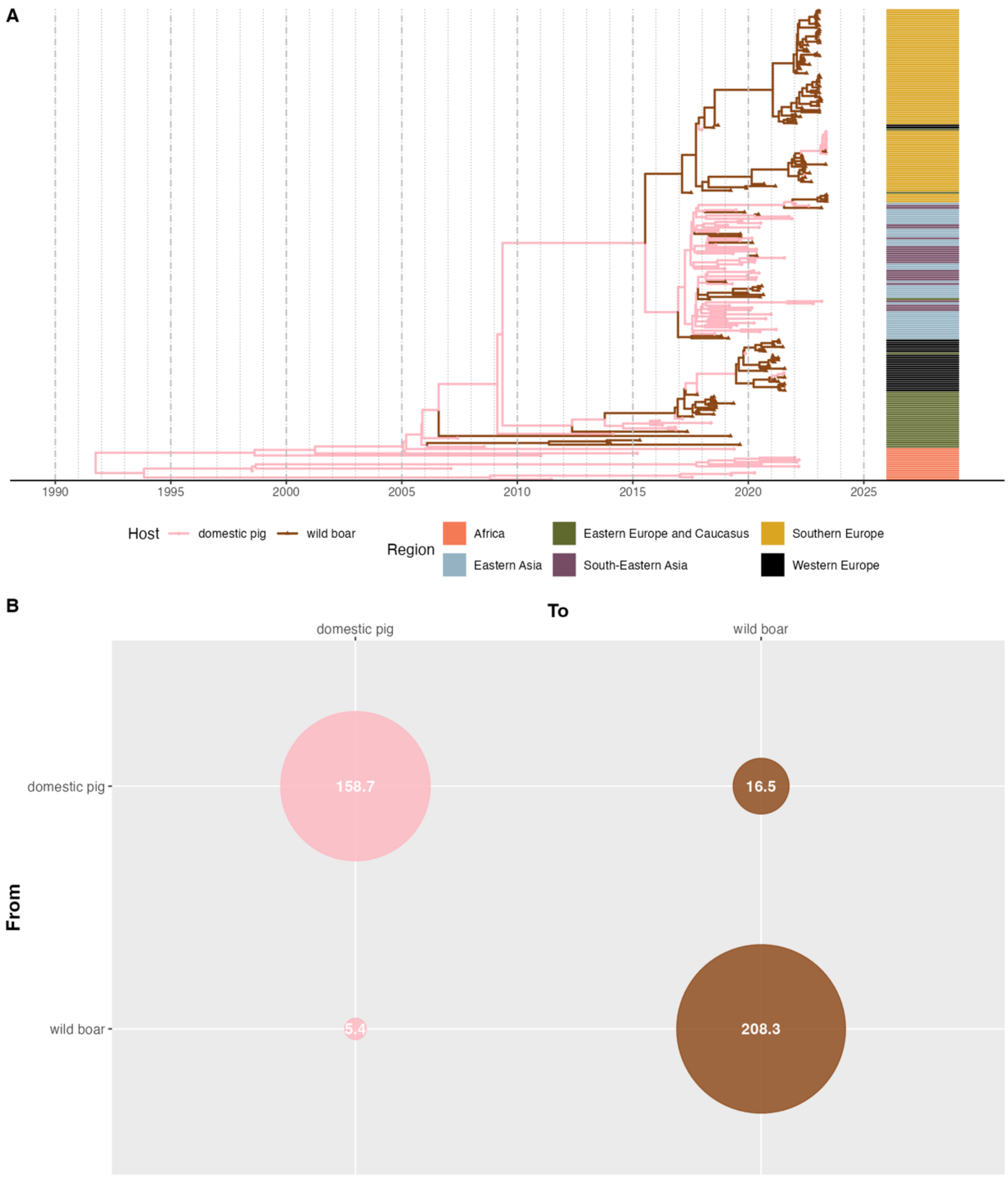


**Figure S5**. Panel A, The Maximum Clade Credibility (MCC) tree of the 217 African Swine fever whole-genome sequences used in the study. Branches are coloured according to the estimated or observed host species (pink for domestic pigs, brown for wild boar or feral pigs). The column on the right report the tips' region (Africa, red; Eastern Asia, blue; South-Eastern Asia, purple; Eastern Europe and Caucasus, green; Southern Europe, yellow; Western Europe, black). Panel B: the estimated transition matrix between domestic pig and wild boar (or feral pigs).

**Movie S1**. The spatially explicit Maximum Clade Credibility tree showing the global spread of ASFV estimated by BEAST in a collection of quarterly snapshots from 1998 to 2023. Black-circled dots represent sequences' (i.e. tips) location (reported or cantered within the smallest available administrative area), small dots represent the estimated internal nodes, arrows represent the tree's branches, and transparent areas correspond to the 80% high-posterior density. Colours show the estimated (for branches or internal nodes) or observed (for tips) region (Africa, red; Eastern Asia, blue; South-Eastern Asia, purple; Eastern Europe and Caucasus, green; Southern Europe, yellow; Western Europe, black). [caption only, file uploaded separately]

| # | Accession number | Original sequence name | Original date | Country | Region |
|---|---|---|---|---|---|
| 1 | FR682468.2 | ASFV Georgia 2007/1\|monopartite | 04/06/2007 | Georgia | Eastern Europe and Caucasus |
| 2 | OP781313.1 | OP781313.1AfricanswinefevervirusisolateMOZ/01/2005,completegenome | 2005 | Mozambique | Africa |
| 3 | OR660695.1 | OR660695.1AfricanswinefevervirusisolateDG_6511_21_,completegenome | 2021 | Serbia | Southern Europe |
| 4 | OR660696.1 | OR660696.1AfricanswinefevervirusisolateDG_6314_19,completegenome | 2019 | Serbia | Southern Europe |
| 5 | OR660698.1 | OR660698.1AfricanswinefevervirusisolateDG_6759_19,completegenome | 2019 | Serbia | Southern Europe |
| 6 | OR660699.1 | OR660699.1Africanswinefevervirusisolate7540/22,completegenome | 2022 | Serbia | Southern Europe |
| 7 | LR722599.1 | ASFV Moldova 2017/1\|monopartite | 2017 | Moldova | Eastern Europe and Caucasus |
| 8 | OR162436.1 | Korea/CW714/2020\|wild boar\|South Korea\|2020 | 2020 | South Korea | Eastern Asia |
| 9 | OP628183.1 | Korea/HC224/2020\|wildboar\|South Korea\|2020 | 2020 | South Korea | Eastern Asia |
| 10 | ON075797.1 | Korea/YC1/2019\|wild boar\|South Korea\|2019 | 2019 | South Korea | Eastern Asia |
| 11 | MT748042.2 | ASFV/Korea/pig/PaJu1/2019\|domestic pig\|II\|South Korea: PaJu\|16-Sep-2019\|A isolate in porcine alveolar macrophage (PAM) | 16-Sep-19 | South Korea | Eastern Asia |
| 12 | ON263123.1 | GZ201801_2\|domestic pig\|II\|China\|serum\|22-Dec-2018 | 22-Dec-18 | China | Eastern Asia |
| 13 | OP467597.1 | ASF-MNG19\|swine\|II\|Mongolia\|2019 | 2019 | Mongolia | Eastern Asia |
| 14 | PP050512.1 | PP050512.1 African swine fever virus isolate 24684_2361/SA/2023_Ita, complete genome | 22/05/2023 | Italy | Southern Europe |
| 15 | PP050513.1 | PP050513.1 African swine fever virus isolate 24685.1_2365/SA/2023_Ita, complete genome | 22/05/2023 | Italy | Southern Europe |
| 16 | PP050515.1 | PP050516.1 African swine fever virus isolate 24689_2369/SA/2023_Ita, complete genome | 22/05/2023 | Italy | Southern Europe |
| 17 | PP050514.1 | PP050514.1 African swine fever virus isolate 24688_2368/SA/2023_Ita, complete genome | 22/05/2023 | Italy | Southern Europe |
| 18 | OP510032.1 | ASFV/Primorsky_19/DP-8235\|Domestic pig\|Russia\|bone marrow\|17-Sep-2019 | 17-Sep-19 | Russia | Eastern Asia |
| 19 | LR722600.1 | ASFV CzechRepublic 2017/1\|monopartite | 17-Jul | Czech Republic | Eastern Europe and Caucasus |
| 20 | LR536725.1 | ASFV Belgium 2018/1\|monopartite | 2018 | Belgium | Western Europe |
| 21 | OM481275.1 | ABTCVSCK_ASF001\|India: Meghalaya\|2020 | 13/05/2020 | India | South-Eastern Asia |
| 22 | OM481276.1 | ABTCVSCK_ASF007\|India: Assam\|2021 | 22/07/2021 | India | South-Eastern Asia |
| 23 | OP605386.1 | 20355/RM/2022_Italy\|Sus scrofa\|Italy: Rome\|spleen\|2022 | 29/04/2022 | Italy | Southern Europe |
| 24 | OR460730.1 | OR460730.1 African swine fever virus isolate 21730_1474/RM/2022_Ita, complete genome | 09/05/2022 | Italy | Southern Europe |
| 25 | OR460735.1 | OR460735.1 African swine fever virus isolate 34616_2119/RM/2022_Ita, complete genome | 18/06/2022 | Italy | Southern Europe |
| *26* | PV833565.1 | PV833565.1 African swine fever virus isolate 21442_1470/RM/2022_Ita, complete genome | 06/05/2022 | Italy | Southern Europe |
| 27 | PP050529.1 | PP050529.1 African swine fever virus isolate 34619_2122/RM/2022_Ita, complete genome | 23/06/2022 | Italy | Southern Europe |
| *28* | PV833566.1 | PV833566.1 African swine fever virus isolate 22283_1480/RM/2022_Ita, complete genome | *12/05/2022* | *Italy* | *Southern Europe* |

| | | | | | |
|---|---|---|---|---|---|
| 29 | PP050528.1 | PP050528.1 African swine fever virus isolate 34611_2114/RM/2022_Ita, complete genome | 12/06/2022 | Italy | Southern Europe |
| 30 | OR460734.1 | OR460734.1 African swine fever virus isolate 34612_2115/RM/2022_Ita, complete genome | 12/06/2022 | Italy | Southern Europe |
| 31 | PP050539.1 | PP050539.1 African swine fever virus isolate 50665.13_2175/RM/2022_Ita, complete genome | 27/08/2022 | Italy | Southern Europe |
| 32 | PP050525.1 | PP050525.1 African swine fever virus isolate 21826_2300/RM/2023_Ita, complete genome | 08/05/2023 | Italy | Southern Europe |
| 33 | PV833567.1 | PV833567.1 African swine fever virus isolate 34597_2126/RM/2022_Ita, complete genome | 09/06/2022 | Italy | Southern Europe |
| 34 | PP050526.1 | PP050526.1 African swine fever virus isolate 34606_2109/RM/2022_Ita, complete genome | 01/06/2022 | Italy | Southern Europe |
| 35 | PP050527.1 | PP050527.1 African swine fever virus isolate 34607_2110/RM/2022_Ita, complete genome | 08/06/2022 | Italy | Southern Europe |
| 36 | OL692743.1 | IND/AS/SD-02/2020\|Domestic Pig\|II\|India\|Apr-2020 | 20-Apr | India | South-Eastern Asia |
| 37 | OL692744.1 | IND/AR/SD-61/2020\|Domestic Pig\|II\|India\|Apr-2020 | 20-Apr | India | South-Eastern Asia |
| 38 | MW465755.1 | VNUA-ASFV-05L1/HaNam/VN/2020\|Sus scrofa\|II\|Viet Nam\|spleen\|2020 | 2020 | Viet Nam | South-Eastern Asia |
| 39 | LC659086.1 | LC659086.1AfricanswinefevervirusAQS-C-1-21DNA,completegenome | Jan-2019/Dec-2020 | China | Eastern Asia |
| 40 | LC659087.1 | LC659087.1AfricanswinefevervirusAQS-C-1-22DNA,completegenome | Jan-2019/Dec-2020 | China | Eastern Asia |
| 41 | OR126359.1 | Pig/Hubei/628/2020\|Domestic pig\|genotype II\|China\|Lymph nodes\|Jun-2020 | 20-Jun | China | Eastern Asia |
| 42 | MW033528.1 | 8\|ASFV-wbShX01\|wild boar\|II\|China\|01-Nov-2019 | 01-Nov-19 | China | Eastern Asia |
| 43 | LC659088.1 | LC659088.1AfricanswinefevervirusAQS-P-20901-1DNA,completegenome | Jan-2019/Dec-2020 | Philippines | South-Eastern Asia |
| 44 | LC659089.1 | LC659089.1AfricanswinefevervirusAQS-P-201202DNA,completegenome | Jan-2019/Dec-2020 | Philippines | South-Eastern Asia |
| 45 | MW306191.1 | ASFV/Primorsky 19/WB-6723\|wild boar\|Russia\|spleen\|28-Aug-2019 | 28-Aug-19 | Russia | Eastern Asia |
| 46 | OP612151.1 | SY-2\|bama mini-pig\|China: Wuhan,Hubei Province\|Oct-2021 | 21-Oct | China | Eastern Asia |
| 47 | OM161110.1 | SY-1\|wild boar\|China\|spleen\|2020-06 | Jun-20 | China | Eastern Asia |
| 48 | MK333181.1 | DB/LN/2018\|dried blood\|2\|China\|Sep-2018 | 18-Sep | China | Eastern Asia |
| 49 | MK333180.1 | Pig/HLJ/2018\|domestic pig\|2\|China\|05-Sep-2018 | 05-Sep-18 | China | Eastern Asia |
| 50 | MH766894.3 | ASFV-SY18\|domestic pig\|II\|China\|spleen\|Jul-2018 | 18-Jul | China | Eastern Asia |
| 51 | MN172368.1 | ASFV/pig/China/CAS19-01/2019\|Sus scrofa scrofa\|p72 II\|China: Zhuhai\|spleen\|02-Jan-2019 | 02-Jan-19 | China | Eastern Asia |
| 52 | OP856591.1 | China/LN/2018/1\|domestic pig\|II\|China\|spleen\|03-Aug-2018 | 03-Aug-18 | China | Eastern Asia |
| 53 | MK128995.1 | China/2018/AnhuiXCGQ\|domestic pig\|II\|China\|02-Sep-2018 | 02-Sep-18 | China | Eastern Asia |
| 54 | MT496893.1 | GZ201801\|domestic pig\|China\|porcine serum\|22-Dec-2018 | 22-Dec-18 | China | Eastern Asia |
| 55 | OR180113.1 | ASFV JS\|swine\|China\|2022 | 2022 | China | Eastern Asia |
| 56 | OQ737679.1 | pig/HuB1/2019\|Sus scrofa\|China\|14-May-2019 | 14-May-19 | China | Eastern Asia |
| 57 | MW791752.1 | ASFV2020-008-B\|Sus scrofa domesticus\|II\|Philippines\|27-Feb-2020 | 27-Feb-20 | Philippines | South-Eastern Asia |

| | | | | | |
|---|---|---|---|---|---|
| 58 | MW791754.1 | ASFV2020-014-B\|Sus scrofa domesticus\|II\|Philippines\|02-May-2020 | 02-May-20 | Philippines | South-Eastern Asia |
| 59 | MW791759.1 | ASFV2020-021-B\|Sus scrofa domesticus\|II\|Philippines\|24-Apr-2020 | 24-Apr-20 | Philippines | South-Eastern Asia |
| 60 | MW791761.1 | ASFV2020-003-B\|Sus scrofa domesticus\|II\|Philippines\|29-Jan-2020 | 29-Jan-20 | Philippines | South-Eastern Asia |
| 61 | MW791757.1 | ASFV2020-019-B\|Sus scrofa domesticus\|II\|Philippines\|18-Jun-2020 | 18-Jun-20 | Philippines | South-Eastern Asia |
| 62 | MW791758.1 | ASFV2020-020-B\|Sus scrofa domesticus\|II\|Philippines\|18-Jun-2020 | 18-Jun-20 | Philippines | South-Eastern Asia |
| 63 | MW791755.1 | ASFV2020-015-B\|Sus scrofa domesticus\|II\|Philippines\|05-May-2020 | 05-May-20 | Philippines | South-Eastern Asia |
| 64 | MW791753.1 | ASFV2020-013-B\|Sus scrofa domesticus\|II\|Philippines\|24-Mar-2020 | 24-Mar-20 | Philippines | South-Eastern Asia |
| 65 | MW791760.1 | ASFV2019-003-B\|Sus scrofa domesticus\|II\|Philippines\|06-Dec-2019 | 06-Dec-19 | Philippines | South-Eastern Asia |
| 66 | MN393476.1 | ASFV Wuhan 2019-1\|domestic pig\|China\|19-Aug-2019 | 19-Aug-19 | China | Eastern Asia |
| 67 | MN393477.1 | ASFV Wuhan 2019-2\|domestic pig\|China\|19-Aug-2019 | 19-Aug-19 | China | Eastern Asia |
| 68 | MK940252.1 | 8\|CN/2019/InnerMongolia-AES01\|domestic wild boar\|II\|China\|19-Feb-2019 | 19-Feb-19 | China | Eastern Asia |
| 69 | OM799941.1 | ASFV/Kaliningrad_17/WB-13869\|wild boar\|Russia\|Bone marrow\|07-Nov-2017 | 07-Nov-17 | Russia | Eastern Europe and Caucasus |
| 70 | OM966715.1 | ASFV/Kaliningrad_18/WB-12524\|wild boar\|Russia\|spleen\|30-Jul-2018 | 30-Jul-18 | Russia | Eastern Europe and Caucasus |
| 71 | OM966720.1 | ASFV/Kaliningrad_18/WB-12516\|wild boar\|Russia\|spleen\|07-Aug-2018 | 07-Aug-18 | Russia | Eastern Europe and Caucasus |
| 72 | OM966718.1 | ASFV/Kaliningrad_18/WB-9766\|wild boar\|Russia\|spleen\|08-Jul-2018 | 08-Jul-18 | Russia | Eastern Europe and Caucasus |
| 73 | OM966721.1 | ASFV/Kaliningrad_18/WB-9734\|wild boar\|Russia\|spleen\|25-Jun-2018 | 25-Jun-18 | Russia | Eastern Europe and Caucasus |
| 74 | OM966719.1 | ASFV/Kaliningrad_19/WB-10168\|wild boar\|Russia\|spleen\|13-May-2019 | 13-May-19 | Russia | Eastern Europe and Caucasus |
| 75 | MT847621.1 | Pol18_28298_O111\|Sus scrofa\|II\|Poland\|2017/2019\|Natalia Mazur-Panasiuk | 2017/2019 | Poland | Eastern Europe and Caucasus |
| 76 | MG939588.1 | Pol17_04461_C210\|Sus scrofa\|Field\|2\|pig alveolar macrophages\|Poland\|spleen\|Jan-2016/Dec-2017\|Natalia Mazur | Jan-2016/Dec-2017 | Poland | Eastern Europe and Caucasus |
| 77 | MG939583.1 | Pol16_20186_o7\|Sus scrofa\|Field\|2\|pig alveolar macrophages\|Poland\|spleen\|Jan-2016/Dec-2017\|Natalia Mazur | Jan-2016/Dec-2017 | Poland | Eastern Europe and Caucasus |
| 78 | MG939585.1 | Pol16_20540_o10\|Sus scrofa\|Field\|2\|pig alveolar macrophages\|Poland\|spleen\|Jan-2016/Dec-2017\|Natalia Mazur | Jan-2016/Dec-2017 | Poland | Eastern Europe and Caucasus |
| 79 | MG939586.1 | Pol16_29413_o23\|Sus scrofa\|Field\|2\|pig alveolar macrophages\|Poland\|spleen\|Jan-2016/Dec-2017\|Natalia Mazur | Jan-2016/Dec-2017 | Poland | Eastern Europe and Caucasus |
| 80 | MG939587.1 | Pol17_03029_C201\|Sus scrofa\|Field\|2\|pig alveolar macrophages\|Poland\|spleen\|Jan-2016/Dec-2017\|Natalia Mazur | Jan-2016/Dec-2017 | Poland | Eastern Europe and Caucasus |
| 81 | MG939589.1 | Pol17_05838_C220\|Sus scrofa\|Field\|2\|pig alveolar macrophages\|Poland\|spleen\|Jan-2016/Dec-2017\|Natalia Mazur | Jan-2016/Dec-2017 | Poland | Eastern Europe and Caucasus |
| 82 | OM966714.1 | ASFV/Kaliningrad_18/WB-12523\|wild boar\|Russia\|spleen\|07-Aug-2018 | 07-Aug-18 | Russia | Eastern Europe and Caucasus |
| 83 | OM966716.1 | ASFV/Kaliningrad_18/WB-9735\|wild boar\|Russia\|spleen\|03-Jul-2018 | 03-Jul-18 | Russia | Eastern Europe and Caucasus |
| 84 | OM966717.1 | ASFV/Kaliningrad_18/WB-9763\|wild boar\|Russia\|bone marrow\|07-Jul-2018 | 07-Jul-18 | Russia | Eastern Europe and Caucasus |
| 85 | MT847620.1 | Pol17_55892_C754\|Sus scrofa\|II\|Poland\|2017/2019\|Natalia Mazur-Panasiuk | 2017/2019 | Poland | Eastern Europe and Caucasus |
| 86 | MT847622.1 | Pol17_31177_O81\|Sus scrofa\|II\|Poland\|2017/2019\|Natalia Mazur-Panasiuk | 2017/2019 | Poland | Eastern Europe and Caucasus |

| | | | | | |
|---|---|---|---|---|---|
| 87 | MT847623.2 | Pol19_53050_C1959/19\|Sus scrofa\|II\|Poland\|2017/2019\|Natalia Mazur-Panasiuk | 2017/2019 | Poland | Eastern Europe and Caucasus |
| 88 | MK543947.1 | Belgium/Etalle/wb/2018\|wild Boar\|Belgium\|10-Sep-2018 | 10-Sep-18 | Belgium | Western Europe |
| 89 | OP781312.1 | OP781312.1AfricanswinefevervirusisolateRSA/08/2019,completegenome | 2019 | South Africa | Africa |
| 90 | MN715134.1 | ASFV_HU_2018\|wild boar\|Hungary\|porcine alveolar macrophages\|24-Apr-2018 | 24-Apr-18 | Hungary | Eastern Europe and Caucasus |
| 91 | MW396979.1 | ASFV/Timor-Leste/2019/1\|domestic pig\|Timor-Leste\|2019 | 2019 | Timor-Leste | South-Eastern Asia |
| 92 | OP781311.1 | OP781311.1AfricanswinefevervirusisolateZIM/2015,completegenome | 2015 | Zimbabwe | Africa |
| 93 | OR159217.1 | S-S-VR-413000-00015\|wildboar\|South Korea\|2020 | 2020 | South Korea | Eastern Asia |
| 94 | OR159219.1 | 20s2287\|S-S-VR-413000-00002\|wildboar\|South Korea\|2020 | 2020 | South Korea | Eastern Asia |
| 95 | OP510033.1 | ASFV/Zabaykali_20/DP-4905\|Domestic pig\|Russia\|blood\|14-Jul-2020 | 14-Jul-20 | Russia | Eastern Asia |
| 96 | OR159218.1 | 20s2881\|S-S-VR-413000-00008\|wildboar\|South Korea\|2020 | 2020 | South Korea | Eastern Asia |
| 97 | MK645909.1 | MK645909.1AfricanswinefevervirusisolateASFV-wbBS01,completegenome | 01-Nov-18 | China | Eastern Asia |
| 98 | ON456300.2 | Yangzhou\|pig\|type II\|China\|primary alveolar macrophages from lymph node\|Nov-2021 | 21-Nov | China | Eastern Asia |
| 99 | MW306190.1 | ASFV/Amur 19/WB-6905\|wild boar\|Russia\|spleen\|29-Aug-2019 | 29-Aug-19 | Russia | Eastern Asia |
| 100 | MZ614662.1 | CADC_HN09\|swine\|China\|whole blood\|2019 | 2019 | China | Eastern Asia |
| 101 | MW306192.1 | ASFV/Ulyanovsk 19/WB-5699\|wild boar\|Russia\|spleen\|21-Aug-2019 | 21-Aug-19 | Russia | Eastern Europe and Caucasus |
| 102 | MK628478.1 | ASFV/LT14/1490\|wild boar\|II\|domestic pig\|Lithuania\|blood\|Jan-2014 | 14-Jan | Lithuania | Eastern Europe and Caucasus |
| 103 | MH681419.1 | ASFV/POL/2015/Podlaskie\|wild boar\|domestic pig erythrocytes\|Poland\|spleen\|2015 | 2015 | Poland | Eastern Europe and Caucasus |
| 104 | OP781310.1 | OP781310.1AfricanswinefevervirusisolateMAL/04/2011,completegenome | 2011 | Malawi | Africa |
| 105 | OR604566.1 | OR604566.1AfricanswinefevervirusisolateK1-1/Liver/Kupang/Indonesia/2023,completegenome | Feb-March 2023 | Indonesia | South-Eastern Asia |
| 106 | MT882025.1 | VN/QP-ASFV1(2019)\|Sow\|Viet Nam\|spleen\|02-May-2019 | 02-May-19 | Viet Nam | South-Eastern Asia |
| 107 | MT872723.1 | VN/HY-ASFV1(2019)\|domestic pig\|Viet Nam\|Feb-2019 | 19-Feb | Viet Nam | South-Eastern Asia |
| 108 | OR660697.1 | OR660697.1AfricanswinefevervirusisolateDG_167_20_2,completegenome | 2020 | Serbia | Southern Europe |
| 109 | OX376250.1 | 2020ASP02805\|monopartite | 16/11/2020 | Germany | Western Europe |
| 110 | OX376258.1 | 2020ASP02894\|monopartite | 26/11/2020 | Germany | Western Europe |
| 111 | OX376251.1 | 2021ASP01919\|monopartite | 19/04/2021 | Germany | Western Europe |
| 112 | OX376254.1 | 2021ASP00484\|monopartite | 21/01/2021 | Germany | Western Europe |
| 113 | OX376255.1 | 2021ASP00902\|monopartite | 18/02/2021 | Germany | Western Europe |
| 114 | OX376260.1 | 2021ASP01917\|monopartite | 19/04/2021 | Germany | Western Europe |
| 115 | OX376272.1 | 2021ASP03740\|monopartite | 29/07/2021 | Germany | Western Europe |

| | | | | | |
|---|---|---|---|---|---|
| 116 | OX376253.1 | 2021ASP01957\|monopartite | 20/04/2021 | Germany | Western Europe |
| 117 | OX376262.1 | 2021ASP00921\|monopartite | 22/02/2021 | Germany | Western Europe |
| 118 | OX376259.1 | 2021ASP02665\|monopartite | 11/05/2021 | Germany | Western Europe |
| 119 | OX376273.1 | 2021ASP03711\|monopartite | 28/07/2021 | Germany | Western Europe |
| 120 | OX376266.1 | 2021ASP03643\|monopartite | 26/07/2021 | Germany | Western Europe |
| 121 | OX376271.1 | 2021ASP03658\|monopartite | 27/07/2021 | Germany | Western Europe |
| 122 | OX376267.1 | 2021ASP03251\|monopartite | 12/07/2021 | Germany | Western Europe |
| 123 | OX376264.1 | 2021ASP03380\|monopartite | 16/07/2021 | Germany | Western Europe |
| 124 | OX376268.1 | 2021ASP03144\|monopartite | 29/06/2021 | Germany | Western Europe |
| 125 | OX376252.1 | 2021ASP00703\|monopartite | 09/02/2021 | Germany | Western Europe |
| 126 | LR899193.1 | ASFV Germany 2020/1\|monopartite | 24/08/2020 | Germany | Western Europe |
| 127 | OX376257.1 | 2021ASP02148\|monopartite | 29/04/2021 | Germany | Western Europe |
| 128 | OX376263.1 | 2021ASP02207\|monopartite | 06/05/2021 | Germany | Western Europe |
| 129 | OX376256.1 | 2020ASP01832\|monopartite | 20/09/2020 | Germany | Western Europe |
| 130 | OX376261.1 | 2020ASP02103\|monopartite | 07/10/2020 | Germany | Western Europe |
| 131 | OX376265.1 | 2021ASP03384\|monopartite | 16/07/2021 | Germany | Western Europe |
| 132 | PP317788.1 | PP317788.1 African swine fever virus isolate 1054_1434/AL/2022_Ita, complete genome | 03/01/2022 | Italy | Southern Europe |
| 133 | PP317811.1 | PP317811.1 African swine fever virus isolate 47169.13_1496/AL/2022_Ita, complete genome | 12/03/2022 | Italy | Southern Europe |
| 134 | OR460741.1 | OR460741.1 African swine fever virus isolate 50665.12_2152/GE/2022_Ita, complete genome | 25/07/2022 | Italy | Southern Europe |
| 135 | OR460740.1 | OR460740.1 African swine fever virus isolate 50665.9_2157/AL/2022_Ita, complete genome | 30/04/2022 | Italy | Southern Europe |
| 136 | OR460737.1 | OR460737.1Africanswinefevervirusisolate50665.4_2159/AL/2022_Ita,completegenome | 28/04/2022 | Italy | Southern Europe |
| 137 | PP025332.1 | PP025332.1Africanswinefevervirusisolate2077_1448/GE/2022_Ita,completegenome | 13/01/2022 | Italy | Southern Europe |
| 138 | OR460733.1 | OR460733.1Africanswinefevervirusisolate47169.14_1497/AL/2022_Ita,completegenome | 02/04/2022 | Italy | Southern Europe |
| 139 | PP317791.1 | PP317791.1 African swine fever virus isolate 22700_2602/AL/2023_Ita, complete genome | 15/01/2023 | Italy | Southern Europe |
| *140* | PV833568.1 | PV833568.1 African swine fever virus isolate 8549_2280/AL/2023_Ita, complete genome | 10/01/2023 | Italy | Southern Europe |
| 141 | PP317807.1 | PP317807.1 African swine fever virus isolate 22700_2642/AL/2023_Ita, complete genome | 02/02/2023 | Italy | Southern Europe |
| 142 | PP317792.1 | PP317792.1 African swine fever virus isolate 22700_2607/AL/2023_Ita, complete genome | 19/01/2023 | Italy | Southern Europe |
| 143 | PP317798.1 | PP317798.1 African swine fever virus isolate 22700_2623/AL/2023_Ita, complete genome | 26/01/2023 | Italy | Southern Europe |
| 144 | PP317819.1 | PP317819.1 African swine fever virus isolate 8549_2267/AL/2022_Ita, complete genome | 30/12/2022 | Italy | Southern Europe |

| | | | | | |
|---|---|---|---|---|---|
| 145 | PP317808.1 | PP317808.1 African swine fever virus isolate 22700_2644/AL/2023_Ita, complete genome | 12/02/2023 | Italy | Southern Europe |
| 146 | PP317800.1 | PP317800.1 African swine fever virus isolate 22700_2625/AL/2023_Ita, complete genome | 29/01/2023 | Italy | Southern Europe |
| 147 | PP317803.1 | PP317803.1 African swine fever virus isolate 22700_2631/AL/2023_Ita, complete genome | 29/01/2023 | Italy | Southern Europe |
| 148 | PP317814.1 | PP317814.1 African swine fever virus isolate 8549_2250/AL/2022_Ita, complete genome | 02/12/2022 | Italy | Southern Europe |
| 149 | PP317801.1 | PP317801.1 African swine fever virus isolate 22700_2627/AL/2023_Ita, complete genome | 27/01/2023 | Italy | Southern Europe |
| 150 | PP025335.1 | PP025335.1 African swine fever virus isolate 8549_2232/GE/2022_Ita, complete genome | 16/12/2022 | Italy | Southern Europe |
| 151 | PP317804.1 | PP317804.1 African swine fever virus isolate 22700_2633/AL/2023_Ita, complete genome | 31/01/2023 | Italy | Southern Europe |
| 152 | PP317820.1 | PP317820.1 African swine fever virus isolate 8549_2269/AL/2022_Ita, complete genome | 30/12/2022 | Italy | Southern Europe |
| 153 | PP317805.1 | PP317805.1 African swine fever virus isolate 22700_2635/AL/2023_Ita, complete genome | 31/01/2023 | Italy | Southern Europe |
| 154 | PP317793.1 | PP317793.1 African swine fever virus isolate 22700_2608/AL/2023_Ita, complete genome | 19/01/2023 | Italy | Southern Europe |
| 155 | ON108571.3 | 2802/AL/2022 Italy\|Sus scrofa\|Italy: Province of Alessandria, Piedmont\|spleen\|2022 | 16/01/2022 | Italy | Southern Europe |
| 156 | PP317817.1 | PP317817.1 African swine fever virus isolate 8549_2260/AL/2022_Ita, complete genome | 22/12/2022 | Italy | Southern Europe |
| 157 | PP317809.1 | PP317809.1 African swine fever virus isolate 22700_2645/AL/2023_Ita, complete genome | 24/02/2023 | Italy | Southern Europe |
| 158 | PP317821.1 | PP317821.1 African swine fever virus isolate 8549_2284/AL/2023_Ita, complete genome | 11/01/2023 | Italy | Southern Europe |
| 159 | PP317810.1 | PP317810.1 African swine fever virus isolate 22700_2646/AL/2023_Ita, complete genome | 24/02/2023 | Italy | Southern Europe |
| 160 | PP317789.1 | PP317789.1 African swine fever virus isolate 22700_2598/AL/2023_Ita, complete genome | 14/01/2023 | Italy | Southern Europe |
| 161 | OR460731.1 | OR460731.1Africanswinefevervirusisolate47169.12_1495/GE/2022_Ita,completegenome | 27/01/2022 | Italy | Southern Europe |
| 162 | PP317790.1 | PP317790.1 African swine fever virus isolate 22700_2600/AL/2023_Ita, complete genome | 14/01/2023 | Italy | Southern Europe |
| 163 | PP025334.1 | PP025334.1 African swine fever virus isolate 50665.15_2154/GE/2022_Ita, complete genome | 24/08/2022 | Italy | Southern Europe |
| 164 | PP025338.1 | PP025338.1 African swine fever virus isolate 8549_2238/GE/2022_Ita, complete genome | 22/12/2022 | Italy | Southern Europe |
| 165 | PP317813.1 | PP317813.1 African swine fever virus isolate 50665.7_2173/AL/2022_Ita, complete genome | 10/07/2022 | Italy | Southern Europe |
| 166 | PP317794.1 | PP317794.1 African swine fever virus isolate 22700_2612/AL/2023_Ita, complete genome | 22/01/2023 | Italy | Southern Europe |
| 167 | OR460736.1 | OR460736.1 African swine fever virus isolate 50665.1_2170/AL/2022_Ita, complete genome | 04/08/2022 | Italy | Southern Europe |
| 168 | PP317812.1 | PP317812.1 African swine fever virus isolate 50665.2_2172/AL/2022_Ita, complete genome | 10/07/2022 | Italy | Southern Europe |
| 169 | OR460738.1 | OR460738.1 African swine fever virus isolate 50665.5_2168/AL/2022_Ita, complete genome | 02/06/2022 | Italy | Southern Europe |
| 170 | PP317816.1 | PP317816.1 African swine fever virus isolate 8549_2256/AL/2022_Ita, complete genome | 17/12/2022 | Italy | Southern Europe |
| *171* | PV833569.1 | PV833569.1 African swine fever virus isolate 50665.3_2163/AL/2022_Ita, complete genome | 23/05/2022 | Italy | Southern Europe |
| 172 | PP317815.1 | PP317815.1 African swine fever virus isolate 8549_2253/AL/2022_Ita, complete genome | 12/12/2022 | Italy | Southern Europe |
| 173 | PP317796.1 | PP317796.1 African swine fever virus isolate 22700_2617/AL/2023_Ita, complete genome | 25/01/2023 | Italy | Southern Europe |

| | | | | | |
|---|---|---|---|---|---|
| 174 | PP317818.1 | PP317818.1 African swine fever virus isolate 8549_2263/AL/2022_Ita, complete genome | 23/12/2022 | Italy | Southern Europe |
| 175 | PP317795.1 | PP317795.1 African swine fever virus isolate 22700_2613/AL/2023_Ita, complete genome | 22/01/2023 | Italy | Southern Europe |
| 176 | PP025337.1 | PP025337.1 African swine fever virus isolate 8549_2235/GE/2022_Ita, complete genome | 20/12/2022 | Italy | Southern Europe |
| 177 | OR460739.1 | OR460739.1 African swine fever virus isolate 50665.8_2167/AL/2022_Ita, complete genome | 26/05/2022 | Italy | Southern Europe |
| 178 | PP025336.1 | PP025336.1 African swine fever virus isolate 8549_2233/SV/2022_Ita, complete genome | 17/12/2022 | Italy | Southern Europe |
| 179 | PP317806.1 | PP317806.1 African swine fever virus isolate 22700_2637/AL/2023_Ita, complete genome | 30/01/2023 | Italy | Southern Europe |
| 180 | PP025333.1 | PP025333.1Africanswinefevervirusisolate47169.11_1494/GE/2022_Ita,completegenome | 21/01/2022 | Italy | Southern Europe |
| 181 | OR460732.1 | OR460732.1Africanswinefevervirusisolate47169.16_1499/GE/2022_Ita,completegenome | 31/03/2022 | Italy | Southern Europe |
| 182 | PP317802.1 | PP317802.1 African swine fever virus isolate 22700_2628/AL/2023_Ita, complete genome | 27/01/2023 | Italy | Southern Europe |
| 183 | PP050517.1 | PP050517.1 African swine fever virus isolate 21896.3_2307/RC/2023_Ita, complete genome | 03/05/2023 | Italy | Southern Europe |
| 184 | PP050518.1 | PP050518.1 African swine fever virus isolate 22489.4_2312/RC/2023_Ita, complete genome | 09/05/2023 | Italy | Southern Europe |
| 185 | PP182138.1 | PP182138.1 African swine fever virus isolate 23276_2329/RC/2023_Ita, complete genome | 11/05/2023 | Italy | Southern Europe |
| 186 | PP050523.1 | PP050523.1 African swine fever virus isolate 23809_2342/RC/2023_Ita, complete genome | 16/05/2023 | Italy | Southern Europe |
| 187 | PP050520.1 | PP050520.1 African swine fever virus isolate 23251_2316/RC/2023_Ita, complete genome | 11/05/2023 | Italy | Southern Europe |
| 188 | PP050521.1 | PP050521.1 African swine fever virus isolate 23260_2325/RC/2023_Ita, complete genome | 11/05/2023 | Italy | Southern Europe |
| 189 | PP182140.1 | PP182140.1 African swine fever virus isolate 23317_2333/RC/2023_Ita, complete genome | 14/05/2023 | Italy | Southern Europe |
| 190 | PP182137.1 | PP182137.1 African swine fever virus isolate 23259_2323/RC/2023_Ita, complete genome | 13/05/2023 | Italy | Southern Europe |
| 191 | PP050522.1 | PP050522.1 African swine fever virus isolate 23324_2335/RC/2023_Ita, complete genome | 14/05/2023 | Italy | Southern Europe |
| 192 | PP050519.1 | PP050519.1 African swine fever virus isolate 23249_2337/RC/2023_Ita, complete genome | 14/05/2023 | Italy | Southern Europe |
| 193 | PP182139.1 | PP182139.1 African swine fever virus isolate 23287_2331/RC/2023_Ita, complete genome | 14/05/2023 | Italy | Southern Europe |
| 194 | PP182136.1 | PP182136.1 African swine fever virus isolate 23254_2321/RC/2023_Ita, complete genome | 13/05/2023 | Italy | Southern Europe |
| 195 | OM105586.1 | LYG18\|Sus scrofa\|China: Lianyungang\|2018-12 | Dec-18 | China | Eastern Asia |
| 196 | MW791756.1 | ASFV2020-018-B\|Sus scrofa domesticus\|II\|Philippines\|24-Feb-2020 | 24-Feb-20 | Philippines | South-Eastern Asia |
| 197 | MW656282.1 | Pig/Heilongjiang/HRB1/2020\|domestic pig\|2\|China\|12-Sep-2020 | 12-Sep-20 | China | Eastern Asia |
| 198 | OK358852.1 | HK_NT_202103\|swine\|Hong Kong\|Mar-2021 | 21-Mar | China | Eastern Asia |
| 199 | MG939584.1 | Pol16_20538_o9\|Sus scrofa\|Field\|2\|pig alveolar macrophages\|Poland\|spleen\|Jan-2016/Dec-2017\|Natalia Mazur | Jan-2016/Dec-2017 | Poland | Eastern Europe and Caucasus |
| 200 | OL622042.1 | wild boar/SNJ/2020\|wild boar\|2\|China\|03-Mar-2020 | 03-Mar-20 | China | Eastern Asia |
| 201 | OQ434234.1 | OQ434234.1AfricanswinefevervirusisolateTAN/01/2011,completegenome | 2011 | Tanzania | Africa |
| 202 | ON409979.1 | TAN/17/Kibaha\|Sus scrofa\|II\|Tanzania\|2017 | 2017 | Tanzania | Africa |

| | | | | | |
|---|---|---|---|---|---|
| 203 | MW856068.1 | 8\|MAL/19/Karonga\|Sus scrofa\|II\|Malawi\|2019\|Lionel Nyabongo and Jean N. Hakizimana | 2019 | Malawi | Africa |
| 204 | ON409983.1 | TAN/20/Morogoro\|Sus scrofa\|II\|Tanzania\|2020 | 2020 | Tanzania | Africa |
| 205 | MT459800.1 | ASFV/Kabardino-Balkaria 19/WB-964\|wildboar\|Russia\|spleen\|26-Mar-2019 | 26-Mar-19 | Russia | Eastern Europe and Caucasus |
| 206 | OP781309.1 | OP781309.1AfricanswinefevervirusisolateMAD/01/1998,completegenome | 1998 | Madagascar | Africa |
| 207 | ON380539.1 | HB03A\|Sus scrofa domesticus\|China\|2020 | 2020 | China | Eastern Asia |
| 208 | ON380540.1 | HB31A\|Sus scrofa domesticus\|China\|2020 | 2020 | China | Eastern Asia |
| 209 | MH910495.1 | Georgia 2008/1\|Georgia\|2008 | 2008 | Georgia | Eastern Europe and Caucasus |
| 210 | OR290104.2 | OR290104.2AfricanswinefevervirusstrainCN/GD/2022,completegenome | 2022 | China | Eastern Asia |
| 211 | NC_044948.1 | Odintsovo_02/14\|Sus scrofa\|II\|Russia\|wild boar\|Feb-2014 | 14-Feb | Russia | Eastern Europe and Caucasus |
| 212 | KP843857.1 | Odintsovo_02/14\|Sus scrofa\|II\|Russia\|wild boar\|Feb-2014 | 14-Feb | Russia | Eastern Europe and Caucasus |
| 213 | ON963982.1 | RC_ON963982.1AfricanswinefevervirusstrainA4,completegenome | 22-Aug | Philippines | South-Eastern Asia |
| 214 | ON400500.1 | YNFN202103\|domestic pig\|II\|China\|porcine serum\|Mar-2021 | 21-Mar | China | Eastern Asia |
| 215 | PP317797.1 | PP317797.1 African swine fever virus isolate 22700_2619/AL/2023_Ita, complete genome | 26/01/2023 | Italy | Southern Europe |
| 216 | PP317799.1 | PP317799.1 African swine fever virus isolate 22700_2624/AL/2023_Ita, complete genome | 26/01/2023 | Italy | Southern Europe |
| 217 | OM105587.1 | JX21\|Sus scrofa\|China: Jiangxi\|2021-03-20 | 20/03/2021 | China | Eastern Asia |
| 218 | OP479889.1 | OP479889.1AfricanswinefevervirusisolateGhana2022-35,completegenome | 04/01/2022 | Ghana | Africa |
| 219 | OP718535.1 | OP718535.1AfricanswinefevervirusisolateGhana2022_62,completegenome | 03/02/2022 | Ghana | Africa |
| 220 | OP718534.1 | OP718534.1AfricanswinefevervirusisolateGhana2022_40,completegenome | 07/03/2022 | Ghana | Africa |
| 221 | OP718533.1 | OP718533.1AfricanswinefevervirusisolateGhana2022-34,completegenome | 11/03/2022 | Ghana | Africa |
| 222 | OR135685.1 | SG/NParks/A-MAM-2023-02-00021\|Sus scrofa\|II\|Singapore\|Feb-2023 | 23-Feb | Singapore | South-Eastern Asia |
| 223 | MN194591.1 | ASFV/Kyiv/2016/131\|Sus scrofa\|Ukraine: Kyiv\|domestic pig farm\|49.381091 N 29.645562 E\|2016-04-11\|State Research Institute of Laboratory Diagnostics and Veterinary Expertise | 11/04/2016 | Ukraine | Eastern Europe and Caucasus |
| 224 | MW521382.1 | HuB20\|domestic swine\|genotype II\|China\|ossa costale\|01-Oct-2020\|Yanyan Zhang | 01-Oct-20 | China | Eastern Asia |
| 225 | OP672342.1 | Nigeria-RV502\|Sus scrofa\|II\|Nigeria: Obio-Akpor, Rivers State\|blood\|24-Jul-2020 | 24-Jul-20 | Nigeria | Africa |
| 226 | OP781308.1 | OP781308.1AfricanswinefevervirusisolateMAU/01/2007,completegenome | 2007 | Mauritius | Africa |
| 227 | LS478113.1 | Estonia 2014\|monopartite | 2014 | Estonia | Eastern Europe and Caucasus |
| 228 | LR881473.1 | Arm/07/CBM/c4\|monopartite | 04/11/2007 | Armenia | Eastern Europe and Caucasus |

**Table S1**. The list of the 228 sequences obtained from GenBank and sampled from the four mainland Italy outbreaks included in the original alignment. Including the Accession number, original sequence name (from GenBank), original date, country, and region.

| # | Substitution model | Clock model | Clock prior (distribution and mean) | Tree prior | MLE (path sampling) | MLE (stone sampling) | MRCA (mean) | Convergence (ESS > 200) |
|---|---|---|---|---|---|---|---|---|
| 1 | HKY +F +I | relax Exp | Normal 1E-5 | Skygrid (27/54) | -259239.4 | -259255 | 1985.39 | NO |
| **2** | **HKY +F +I +G** | **relax Exp** | **Normal 1E-5** | **Skygrid (27/54)** | **-259241.1** | **-259255.2** | **1993.88** | **YES** |
| 3 | GTR +F +I +G | relax Exp | Normal 1E-5 | Skygrid (27/54) | -259278.1 | -259292.8 | 1993.30 | PARTIAL (6 parameters 150<200) |
| 4 | TN93 +F +I +G | relax Exp | Normal 1E-5 | Skygrid (27/54) | -259269.1 | -259284.3 | 1993.00 | YES |
| 5 | HKY +F +I +G | strict | Normal 1E-5 | Skygrid (27/54) | -259381.0 | -259395.1 | 1991.30 | YES |
| 6 | HKY +F +I +G | relax Logn | Normal 1E-5 | Skygrid (27/54) | -259279.0 | -259293.9 | 1990.97 | PARTIAL (8 parameters 100<200) |
| 7 | HKY +F +I +G | random local | Normal 1E-5 | Skygrid (27/54) | -259291.4 | 259305.1 | 1994.19 | NO |
| 8 | HKY +F +I +G | fixed local | Normal 1E-5 | Skygrid (27/54) | -259346.8 | -259360.1 | 1991.37 | YES |
| 9 | HKY +F +I +G | relax Exp | Normal 1E-5 | Constant population | -259318.8 | -259334.5 | 1987.07 | YES |
| 10 | HKY +F +I +G | relax Exp | Normal 1E-5 | Birth-death model | NA | NA | NA | FAILED |
| 11 | HKY +F +I +G | relax Exp | Normal 1E-5 | Expansion growth | -259311.6 | -259327.8 | 1990.07 | YES |
| 12 | HKY +F +I +G | relax Exp | Normal 1E-5 | Exponential growth | -259294.9 | -259310.3 | 1986.74 | YES |
| 13 | HKY +F +I +G | relax Exp | Normal 1E-5 | Logistic growth | NA | NA | NA | FAILED |
| 14 | HKY +F +I +G | relax Exp | Normal 1E-5 | Hamiltonian SkyGrid (27/54) | -259288.4 | -259303.1 | 1993.26 | NO |
| 15 | HKY +F +I +G | relax Exp | Normal 1E-5 | Skygrid (30/60) | -259266.5 | -259281.5 | 1994.07 | PARTIAL (2 parameters 140<200) |
| 16 | HKY +F +I +G | relax Exp | Normal 1E-5 | Skygrid (25/50) | -259256.6 | -259271.2 | 1993.08 | PARTIAL (2 parameters 150<200) |
| 17 | HKY +F +I +G | relax Exp | Uniform 1E-5 | Skygrid (27/54) | -259264.6 | -259279.9 | 1994.10 | PARTIAL (1 parameters 150<200) |
| 18 | HKY +F +I +G | relax Exp | Exponential 1E-5 | Skygrid (27/54) | -259284.8 | -259299.5 | 1993.07 | PARTIAL (2 parameters 190<200) |
| 19 | HKY +F +I +G | relax Exp | Lognormal 1E-5 | Skygrid (27/54) | -259307.8 | -259324.2 | 1995.70 | YES |
| 20 | HKY +F +I +G | relax Exp | Normal 1E-5, SD 1E-4 | Skygrid (27/54) | -259264.4 | -259278.9 | 1985.09 | PARTIAL (1 parameters 140<200) |
| 21 | TN93 +F +G | strict | Normal clock 1E-5 | Constant population | -259431.4 | -259446.8 | 1991.47 | YES |
| 22 | GTR +F +G | relax Logn | Normal clock 1E-5 | Constant population | -259293.7 | -259309.0 | 1985.09 | PARTIAL (7 parameters 100<200) |
| 23 | GTR+F+G | relax Logn | Normal clock 1E-5 | SkyGrid (27/54) | -259443.6 | -259456.0 | 1992.54 | PARTIAL (11 parameters 100<200, 2<100) |

**Table S2**. The hierarchical *BEAST* model selection results. The final model selected is indicated in bold. In models #21, #22 and #23 we tested the same parameters as in, respectively, Forth et al.[13] and Zhang et al.[5], and Gámbaro et al.[14]. For the SkyGrid model, the two number in parenthesis correspond, respectively, to the time at last transitions and the number of bins.

### References


[1] D.J. Winter, *rentrez: An R package for the NCBI eUtils API*, The R Journal 9 (2017), pp. 520–526.

[2] K. Kameyama, T. Kitamura, K. Okadera, M. Ikezawa, K. Masujin and T. Kokuho, *Usability of Immortalized Porcine Kidney Macrophage Cultures for the Isolation of ASFV without Affecting Virulence*, Viruses 14 (2022), pp. 1794.

[3] *UNSD — Methodology*. Available at https://unstats.un.org/unsd/methodology/m49/#ftn13.

[4] B.Q. Minh, H.A. Schmidt, O. Chernomor, D. Schrempf, M.D. Woodhams, A. Von Haeseler et al., *IQ-TREE 2: New Models and Efficient Methods for Phylogenetic Inference in the Genomic Era*, Molecular Biology and Evolution 37 (2020), pp. 1530–1534.

[5] Y. Zhang, Q. Wang, Z. Zhu, S. Wang, S. Tu, Y. Zhang et al., *Tracing the Origin of Genotype II African Swine Fever Virus in China by Genomic Epidemiology Analysis*, Transboundary and Emerging Diseases 2023 (2023), pp. 4820809.

[6] A. Rambaut, T.T. Lam, L.M. Carvalho and O.G. Pybus, *Exploring the temporal structure of heterochronous sequences using TempEst (formerly Path-O-Gen)*, Virus Evolution 2 (2016), pp. 1–7.

[7] *aglucaci/OpenRDP: An open-source re-implementation of the RDP4 recombination detection program*. Available at https://github.com/aglucaci/OpenRDP.

[8] V. Hill and G. Baele, *Bayesian Estimation of Past Population Dynamics in BEAST 1.10 Using the Skygrid Coalescent Model*, Molecular Biology and Evolution 36 (2019), pp. 2620–2628.

[9] M.S. Gill, P. Lemey, N.R. Faria, A. Rambaut, B. Shapiro and M.A. Suchard, *Improving bayesian population dynamics inference: A coalescent-based model for multiple loci*, Molecular Biology and Evolution 30 (2013), pp. 713–724.

[10] A. Rambaut, A.J. Drummond, D. Xie, G. Baele and M.A. Suchard, *Posterior summarization in Bayesian phylogenetics using Tracer 1.7*, Systematic Biology 67 (2018), pp. 901–904.

[11] G. Baele, P. Lemey, T. Bedford, A. Rambaut, M.A. Suchard and A.V. Alekseyenko, *Improving the Accuracy of Demographic and Molecular Clock Model Comparison While Accommodating Phylogenetic Uncertainty*, Molecular Biology and Evolution 29 (2012), pp. 2157–2167.

[12] H. Zhang, S. Zhao, H. Zhang, Z. Qin, H. Shan and X. Cai, *Vaccines for African swine fever: an update*, Front. Microbiol. 14 (2023).

[13] J.H. Forth, L.F. Forth, S. Lycett, L. Bell-Sakyi, G.M. Keil, S. Blome et al., *Identification of African swine fever virus-like elements in the soft tick genome provides insights into the virus' evolution*, BMC Biology 18 (2020), pp. 136.

[14] F. Gámbaro, L.C. Goatley, T.J. Foster, C. Tennakoon, G.L. Freimanis, S. Van Borm et al., *Exploiting Viral DNA Genomes to Explore the Dispersal History of African Swine Fever Genotype II Lineages in Europe*, Genome Biology and Evolution 17 (2025), pp. evaf102.
