## Supplementary figures and images for "A phylogenetic contribution to understanding the panzootic spread of African swine fever: from the global to the local scale"

### Movie S1

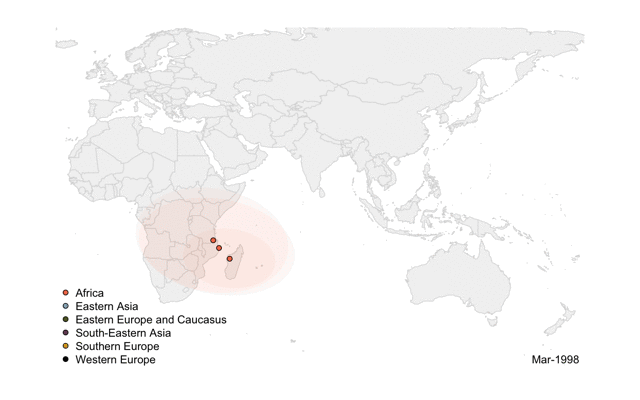
